# The bitter end: T2R bitter receptor agonists elevate nuclear calcium and induce apoptosis in non-ciliated airway epithelial cells

**DOI:** 10.1101/2021.05.16.444376

**Authors:** Derek B. McMahon, Li Eon Kuek, Madeline E. Johnson, Paige O. Johnson, Rachel L.J. Horn, Ryan M. Carey, Nithin D. Adappa, James N. Palmer, Robert J. Lee

## Abstract

Bitter taste receptors (T2Rs) localize to airway motile cilia and initiate innate immune responses in retaliation to bacterial quorum sensing molecules (acyl-homoserine lactones and quinolones). Activation of T2Rs leads to calcium-driven NO production that increases cilia beating and directly kills bacteria. Several airway diseases, including chronic rhinosinusitis, COPD, and cystic fibrosis, are characterized by epithelial remodeling, including loss of motile cilia and/or squamous metaplasia. To understand the function of T2Rs within the altered landscape of airway disease, we studied T2Rs in non-ciliated airway cell lines and primary cells de-differentiated to a squamous phenotype. In differentiated cells, T2Rs localize to cilia, however in de-differentiated, non-ciliated cells they localize to the nucleus. Cilia and nuclear import utilize many shared proteins, thus in the absence of motile cilia some T2Rs may target to the nucleus. T2R agonists selectively elevated both nuclear and mitochondrial calcium through a G-protein-coupled receptor, phospholipase C, and InsP_3_ receptor-dependent mechanism. Additionally, T2R agonists decreased nuclear cAMP, increased nitric oxide, and increased cGMP, consistent with T2R signaling. Furthermore, exposure to T2R agonists led to nuclear calcium-induced mitochondrial depolarization and caspase activation. T2R agonists induced apoptosis in primary bronchial and nasal cells differentiated at air-liquid interface but then induced to a squamous phenotype by apical submersion. Air-exposed well-differentiated cells did not die. This T2R-induced apoptosis may be a last-resort defense against infection, possibly when bacteria have breached the epithelial barrier and reach non-ciliated cells below. However, it may also increase susceptibility of de-differentiated or remodeled epithelia to damage by bacterial metabolites. Moreover, the T2R-activated apoptosis pathway occurs in airway cancer cells. T2Rs may thus contribute to microbiome-tumor cell crosstalk in airway cancers. T2R agonists may also be useful topical therapeutics (e.g., delivered by nasal rinse or nebulizer) for activating airway cancer cell apoptosis without killing surrounding differentiated tissue.

## Introduction

Motile cilia defend the airways. Pathogens get trapped in airway mucus and motile cilia drive their clearance toward the oropharynx for swallowing [1]. Motile cilia are also immune sensors [2]. Cilia contain bitter taste receptors (T2Rs), G protein-coupled receptors (GPCRs) originally identified on the tongue which also detect bacterial N-acylhomoserine lactones and quinolones [3-5]. T2R isoforms 4, 14, 16, and 38, are in human nasal [3, 6] and bronchial [7] cilia. Cilia T2Rs initiate Ca^2+^-triggered nitric oxide (NO) production to increase ciliary beating via cGMP and protein kinase G [3, 5]. NO also damages cell walls and DNA of bacteria [8] and inhibits replication of viruses [9], including SARS-COV-1 and -2 [10, 11].

Clinical data support importance of ciliated cell T2Rs in chronic rhinosinusitis (CRS) [1]. Patients with polymorphisms that render the cilia-localized T2R38 isoform non-functional are at higher risk of CRS, are more likely to require sinus surgery, and may have poorer outcomes after surgery [5]. We hypothesize that pathologies resulting in cilia loss or airway remodeling may impair the T2R pathway and have detrimental effects on innate immunity.

Several airway diseases share phenotypes of acquired cilia defects or loss of cilia due to squamous metaplasia or other inflammatory remodeling, including CRS [1], chronic obstructive pulmonary disease (COPD) [12, 13], and cystic fibrosis (CF) [14]. Loss of cilia occurs with viral or bacterial infection [2], type-2 inflammation-driven remodeling [15], or smoking [16]. Elucidating how T2R signaling changes as the airway epithelium changes is required to understand airway pathophysiology.

We studied how T2R signaling changes when ciliated cells are de-differentiated or replaced with squamous cells in airway disease. We used airway cell lines as well as primary nasal and bronchial cells. While squamous epithelial cells still express T2Rs, altered intracellular localization of T2R-induced Ca^2+^ responses, and possibly the T2Rs themselves, contributes to activation of alternative apoptotic signaling involving Ca^2+^ signaling from the nucleus to mitochondria.

## Materials and Methods

All reagents are shown in **Supplementary Table S1**. More detailed methods are in **Supplementary Material**.

### Human primary cell culture

Primary sinonasal culture was carried out as described [6, 17]. Institutional review board approval (#800614) and written informed consent was obtained. Tissue was collected from patients ≥18 years of age undergoing surgery for sinonasal disease or trans-nasal approaches to the skull base. Primary cells were obtained through dissociation (1.4 mg/ml pronase; 0.1 mg/ml DNase; 1 hour; 37°C) [3]. Primary normal human bronchial (HBE) cells were from Lonza (Walkerville, MD) cultured in PneumaCult (Stemcell Technologies) plus penicillin/streptomycin. Immortalization with BMI-1 (indicated for some experiments) was carried out as in [18]. For deciliation, (HBE) air liquid interface (ALI) cultures were differentiated for 3 weeks and then either exposed to air or apical submersion for 4 days [17]. Age-matched cultures were compared. *TAS2R38* genotyping was previously described [3].

### Cell line culture

Cell culture was as described [6, 17] in Minimal Essential Media with Earl’s salts (Gibco; Gaithersburg, MD USA) plus 10% FBS and 1% penicillin/streptomycin (Gibco). RPMI 2650 (nasal squamous carcinoma), BEAS-2B (adenovirus 12-SV40 hybrid immortalized bronchial), A549 (alveolar type II-like carcinoma), HEK293T, NCI-H292 (lung mucoepidermoid carcinoma expressing T2R14 [19]), and Caco-2 (colorectal adenocarcinoma) cells were from ATCC (Manassas, VA USA). 16HBE (SV-40 immortalized bronchial) cells [20] were from D. Gruenert (University of California San Francisco, San Francisco, CA USA). Transfections used Lipofectamine 3000 (ThermoFisher Scientific, Waltham MA). After transfection with shRNA plasmids for either T2R8, T2R10, T2R14, or scramble shRNA for stable expression, Beas2B cells were supplemented with 1 μg/mL puromycin for 1 week and then maintained in 0.2 μg/mL puromycin.

### Live Cell Imaging

Unless noted, imaging was as described [6, 17]. For Ca^2+^, cells were loaded with 5 μM Fura-2-AM or Fluo-8-AM for 1 hour in HEPES-buffered Hank’s Balanced Salt Solution (HBSS) at room temperature in the dark. Cells were loaded with 10 μM DAF-FM diacetate for 1.5 hours. Fura-2 was imaged using an Olympus IX-83 microscope (20x 0.75 NA objective), fluorescence xenon lamp with excitation and emission filter wheels (Sutter Instruments, Novato, CA USA), Orca Flash 4.0 sCMOS camera (Hamamatsu, Tokyo, Japan) and MetaFluor (Molecular Devices, Sunnyvale, CA USA) using a Fura-2 filters (79002-ET, Chroma, Rockingham, VT USA). Imaging of Fluo-8 or DAF-FM used a FITC filter set (49002-ET, Chroma). For nuclear Ca^2+^ or cAMP, plasmids for G-GECO, R-GECO-nls, Flamindo2 or nls-Flamindo2 were transfected 48 hours prior to imaging. Images were taken with FITC or TRITC filters. Annexin V-FITC was imaged at 10x (0.4 NA).

### Proliferation, Mitochondrial Membrane Potential, and Apoptosis Measurements

XTT was added to sub-confluent cells immediately before absorbance measurements at 475nm (specific absorbance) and 660nm (reference) with Spark 10M plate reader (Tecan; Männedorf, Switzerland). JC-1 dye was added 10 min prior to recording (ex.488/em.535 and em.590) while CellEvent Caspase 3/7 (ThermoFisher) was added immediately prior to recording (ex.495/em.540).

### Immunofluorescence

Cultures were fixed (4% paraformaldehyde, 20 min) followed by incubation in phosphate saline buffer containing 5% normal donkey serum, 1% bovine serum albumen, 0.2% saponin, and 0.1% Triton X-100 for 45 min. Cultures were incubated 1:100 dilutions of T2R or α-gustducin antibodies at 4°C overnight, then AlexaFluor-labeled donkey anti-mouse or anti-rabbit (1:1000) at 4°C for 1 hour. Images were taken on using 60x objective (1.4 NA oil). MitoTracker Deep Red FM was used at 10 nM for 15 min. Co-staining of T2R14 and T2R38 used AlexaFluor 488 and 546 Zenon antibody labeling kits.

### Biochemistry

For endogenous T2Rs, cells were lysed and run on a NuPage 4-12% Bis-Tris gel, transferred to nitrocellulose, then blocked in 5% milk in 50 mM Tris, 150 mM NaCl, and 0.025% Tween-20 (Tris-Tween) for 1 hour. Primary antibody (1:1000) in Tris-Tween with 5% BSA for 1.5 hours. Goat anti-rabbit or anti-mouse IgG-horseradish peroxidase secondary antibodies (1:5000) for 1 hour. Blots were visualized with Clarity ECL on an imager with Image Lab Software (BioRad). Nuclei were isolated using the REAP method [21] then either used for biochemistry or fixed on slides.

### Quantitative PCR (qPCR)

Subconfluent cultures were resuspended in TRIzol (ThermoFisher Scientific). Purified RNA (Direct-zol RNA kit; Zymo Research) was transcribed to cDNA via High-Capacity cDNA RT Kit (ThermoFisher Scientific). Taqman Q-PCR probes were used in a QuantStudio 5 Real-Time PCR System (ThermoFisher Scientific).

### Data analysis and statistics

T-tests (two comparisons only) and one-way ANOVA (>2 comparisons) were calculated using GraphPad PRISM with appropriate post-tests. In all figures, *p* < 0.05 (*), *p* < 0.01 (**), *p* < 0.001 (***).

## Results

### Bitterants regulate Ca^2+^_nuc_

We examined bitter agonist (bitterant)-induced intracellular Ca^2+^ (Ca^2+^ _i_) release in primary human bronchial epithelial (HBE) cells grown in submersion to model non-ciliated squamous basal cells [22] as well as non-ciliated bronchial epithelial lines, including viral-immortalized Beas-2B [23] and 16HBE14o-(16HBE) [20] and cancer-derived A549 and RPMI 2650 cells. T2R agonists diphenhydramine (DPD), flufenamic acid (FFA), and denatonium benzoate all induced Ca^2+^_i_ release in these non-ciliated airway cells (**Supplementary Fig. S1**) and T2R expression was observed (**Supplementary Fig. S2**). Cognate T2Rs for agonists used are in **Table S2**. T2R agonist-induced Ca^2+^_i_ appeared most intense in the nucleus of airway (**Fig. 1A, Supplementary Fig. S3-S5**) and non-airway Caco-2 cells (**Figure S6**). Other GPCRs elicited more global Ca^2+^_i_ responses (**Supplementary Fig. S2, S4**). We saw similar bitterant-induced nuclear Ca^2+^_i_ in primary nasal cells from turbinate brushing cultured in submersion (**Supplementary Fig. S7**).

**Fig. 1.**
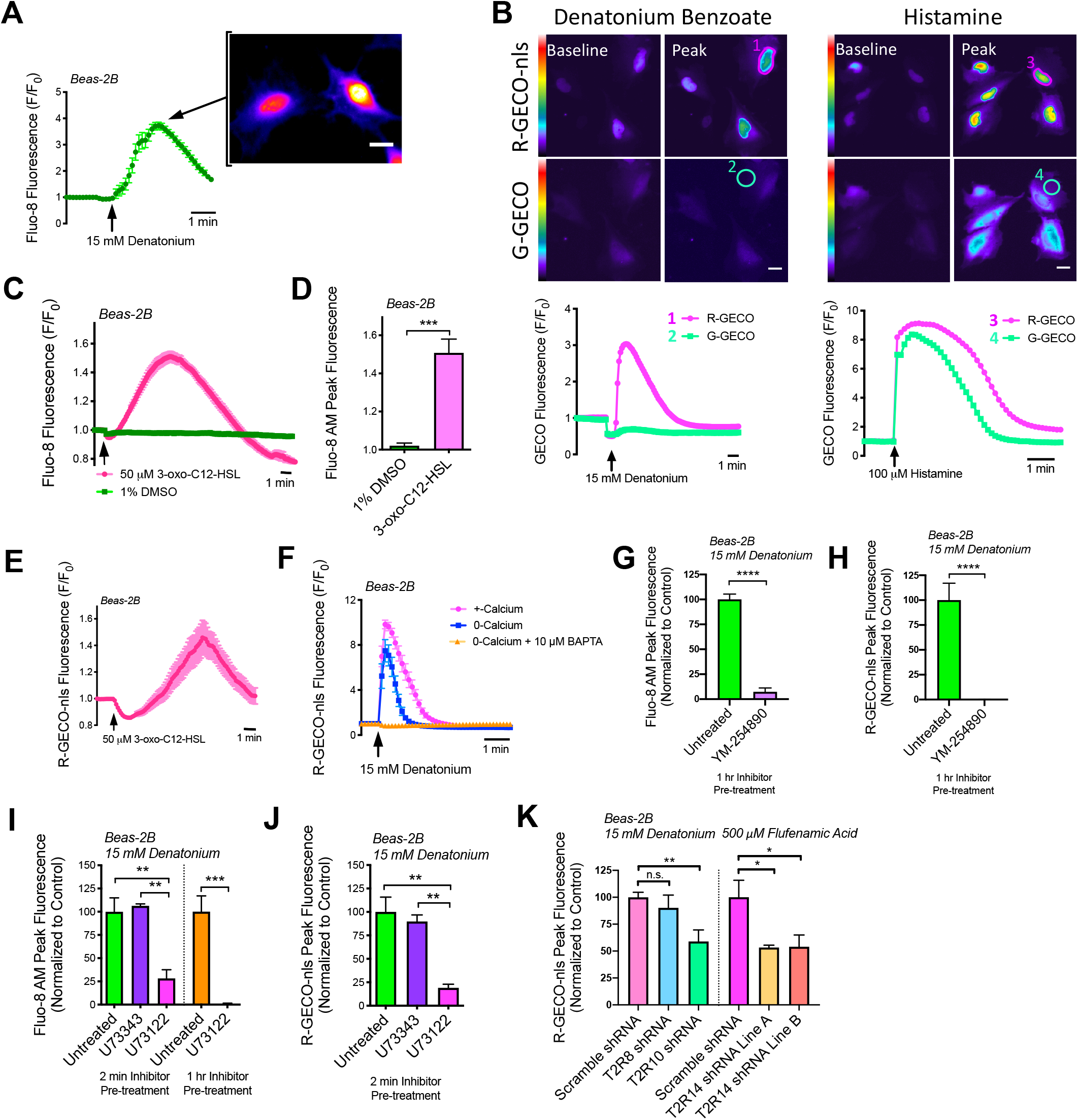
Denatonium-induced T2R-mediated Ca^2+^ release is strongly localized to the nucleus in Beas-2B cells. **A** Beas-2B cells, a bronchial line with squamous phenotype in the presence of FBS [23], were loaded with Ca^2+^ dye Fluo-8. Fluo-8 fluorescence trace and representative image during stimulation with denatonium, showing nuclear Ca^2+^ release. Scale bar = 10 µm. **B** Representative traces showing Beas-2B cells co-expressing G-GECO (Ca^2+^_i_) or R-GECO-nls Ca^2+^_nuc_) and preferential activation of Ca^2+^_nuc_ with denatonium. Scale bar = 10 µm. **C-E** Traces and bar graph showing Ca2+I (Fluo-8; *C-D*) and Ca^2+^_nuc_ (R-GECO-nls; *E*) with quorum sensing molecule and T2R agonist 3-oxo-C12-HSL. **F** Ca^2+^_nuc_ (R-GECO-nls) responses in cells ± 10 μM BAPTA-AM for 1hr ± extracellular calcium. 0-Ca^2+^_o_ buffer contained no added Ca^2+^ plus 2 mM EGTA to chelate trace Ca^2+^. **G-H** Inhibition of denatonium-induced Fluo-8 and R-GECO-nls responses with YM-254890 (1 uM, 1 hr pre-treatment). **I-J** Bar graphs showing inhibition of 15 mM denatonium-induced Ca^2+^_i_ and Ca^2+^_nuc_ with PLC inhibitor U73122 (1 µM; either 2 min or 1 hr pretreatment) but not inactive control analogue U73343. **K** Beas-2B’s stably expressing shRNA targeting either T2R10 or 14, but not T2R8 have reduced bitterant-induced Ca^2+^_nuc_ signaling. All traces are representative experiments from ≥3 independent experiments. Bar graphs show mean ± SEM from ≥3 experiments; significance by T-test (D,G,H) or 1-way ANOVA with Tukey’s posttest (I-K), \*\**P*<0.01, \*\*\**P*<0.001, \*\*\*\**P*<0.0001.

To directly investigate if bitterants elevate nuclear Ca^2+^ (Ca^2+^_nuc_), Beas-2Bs were co-transfected with genetically-encoded Ca^2+^ biosensors: green (G)-GECO and red (R)-GECO-nls [24] to differentiate between global Ca^2+^_i_ and Ca^2+^_nuc_. Denatonium benzoate increased R-GECO-nls more than G-GECO fluorescence (**Fig. 1B**) suggesting T2Rs preferentially increase Ca^2+^_nuc_. In contrast, histamine elevated both similarly, (**Fig. 1B**). Ca^2+^_nuc_ also increased in A549s, RPMI2650s, and submerged primary nasal cells from 3 patients in response to bitterants (**Supplementary Fig. S7-S8**). *Pseudomonas aeruginosa* 3-oxo-C12-HSL activates multiple T2Rs [4, 5]; 50 μM 3-oxo-C12-HSL activated both Ca^2+^_i_ (Fluo-8) and Ca^2+^_nuc_ (R-GECO-nls) in Beas-2Bs (**Fig. 1C-E**). Together, these results demonstrate the novel finding that diverse T2R agonists specifically elevate Ca^2+^_nuc_. Although the cells lack cilia, the T2Rs nonetheless are functional.

We utilized a protocol to permeabilize plasma membrane but not intracellular organelles, previously used to study ER Ca^2+^ release [25]. In permeabilized A549s, quinine still increased Ca^2+^_nuc_ (**Fig. S9**), suggesting Ca^2+^_nuc_ originates from T2R signaling on intracellular membranes. In Beas-2Bs, Ca^2+^_nuc_ was slightly reduced but still intact with 0-Ca^2+^_o_ (no added Ca^2+^ plus 2 mM EGTA) buffer but fully eliminated when cells were preloaded with Ca^2+^ chelator BAPTA (**Fig. 1F**). Thus, T2R-induced Ca^2+^_nuc_ originates largely from internal Ca^2+^ stores, supported by an ER-localized Ca^2+^ biosensor (**Fig. S10**).

Denatonium benzoate activates ∼8 T2Rs, while sodium benzoate activates only 2 with weak affinity [26] (**Table S2**). Denatonium benzoate activated Ca^2+^_nuc_ while equimolar sodium benzoate did not (**Supplementary Fig. S11**). Thus, Ca^2+^_nuc_ is not due to osmolarity or pH. To confirm GPCR involvement, broad-range G protein inhibitor YM-254890 [27] or phospholipase C (PLC) inhibitor U73122 blocked both Ca^2+^_i_ and Ca^2+^_nuc_ (**Fig. 1G-J**). T2R10 or T2R14 shRNAs also reduced Ca^2+^_nuc_ responses to denatonium benzoate (T2R10 agonist) or FFA (T2R14 agonist), respectively (**Fig. 1K**). T2R GPCR signaling is thus responsible for bitterant-induced Ca^2+^_nuc_.

HEK293Ts are often used as model for T2R expression [6, 26]. However, a recent study suggested expression of endogenous T2R14 in HEK293Ts [28]. DPD, which activates T2R14, increased Ca^2+^_nuc_ in HEK293Ts (**Supplementary Fig. S12A-B**). HEK’s express a several T2Rs by qPCR (**Supplementary Fig. S12C**), and a *TAS2R14* promoter GFP reporter revealed bright GFP fluorescence (**Supplementary Fig. S12D**). Staining of HEK293Ts with two different T2R14 antibodies was plasma membrane localized at cell-cell contact points but also partly nuclear (**Supplementary Fig. S12E**). Thus, However, T2R elevation of Ca^2+^_nuc_ may have implications in many cell types and caution should be exercised with HEK’s as a model for T2R expression.

### Bitterants regulate cAMP_nuc_

T2Rs can signal through Gα-gustducin [5] or Gα_i_ [29] to lower cAMP. T2R signaling in ciliated cells is gustducin-independent [3], but nonetheless decreases cAMP [5]. To assess changes nuclear cAMP (cAMP_nuc_), we used cAMP biosensor Flamindo2 and nls-Flamindo2 [30]. T2R agonist diphenidol decreased global intracellular cAMP in Beas-2Bs (**Fig. 2A**). Diphenidol and quinine both decreased cAMP_nuc_ concomitant with increasing Ca^2+^_nuc_ (**Fig. 2B-D**). It remains to be determined if this requires gustducin or a Gα_i_. However, over expression of Wt gustducin reduced Ca^2+^_nuc_ by >50%, while non-functional gustducin had no effect (**Fig. 2E**). There may be partial coupling of Gα_q_ to T2Rs; competing away Gα_q_ with gustducin may lower Ca^2+^ responses. Many GPCRs are G protein promiscuous [31]. Nonetheless, the effects of overexpressing functional gustducin supports T2R involvement.

**Fig. 2.**
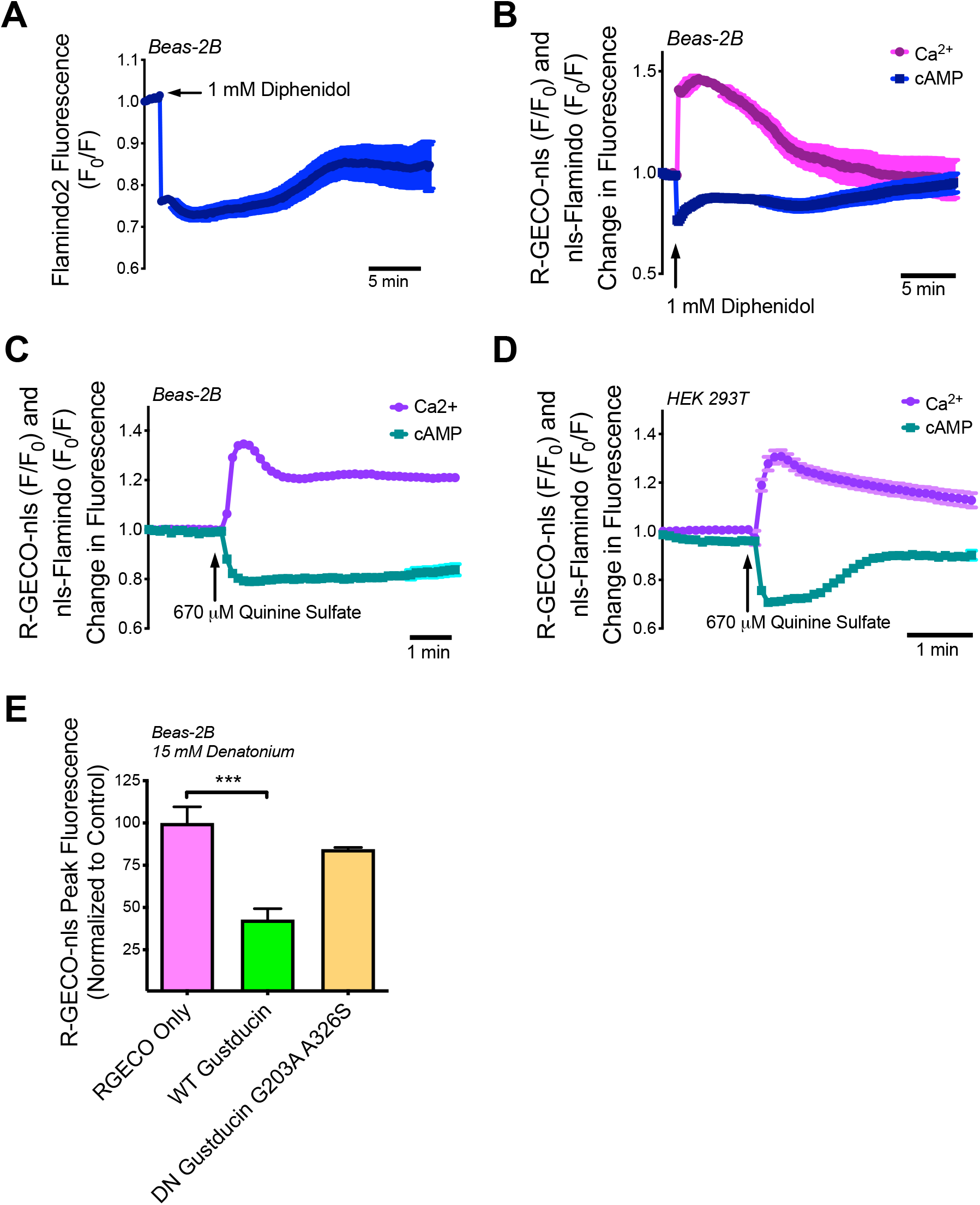
T2R agonists alter cAMP_nuc_ concomitantly with Ca^2+^_nuc_. **A-B** Traces showing cAMP_i_(*A*) and cAMP_nuc_ (*B*) decrease and Ca^2+^_nuc_ increase (*C*) in response to T2R agonist diphenidol in Beas-2B cells. **C-D** Traces showing cAMP_nuc_ and Ca^2+^_nuc_ in response to T2R agonist quinine Beas-2B (*C*) and HEK 239T (*D*) cells. **E** Bar graph showing peak Ca^2+^_nuc_ in Beas-2B cells transfected with WT or non-functional (G203A A326S; [69]) α-gustducin. Traces represent 1 experiment from a minimum of 3 experiments. Bar graph represents mean ± SEM from 3 experiments; significance determined by 1-way ANOVA using Bonferroni post hoc test, \*\*\**P*<0.001.

### Bitterants regulate NO production

T2Rs in differentiated ciliated cells produce NO downstream of Ca^2+^ via endothelial nitric oxide synthase (eNOS) [3, 6]. Both FFA and denatonium benzoate, but not sodium benzoate, increased DAF-FM (NO indicator dye) fluorescence in 16HBEs that was reduced by NO scavenger cPTIO (**Fig. 3A-B**) or NOS inhibitor L-NAME (**Fig. 3C-D**). GENIe cGMP biosensor showed cGMP increases with FFA and denatonium benzoate (**Fig. 3E-H**). Thus, the T2R Ca^2+^ response, even though nuclear, can still increase NO/cGMP. Pretreatment with L-NAME did not alter Fluo-8 responses (**Fig. 3I**), suggesting the NO is downstream of Ca^2+^.

**Fig. 3.**
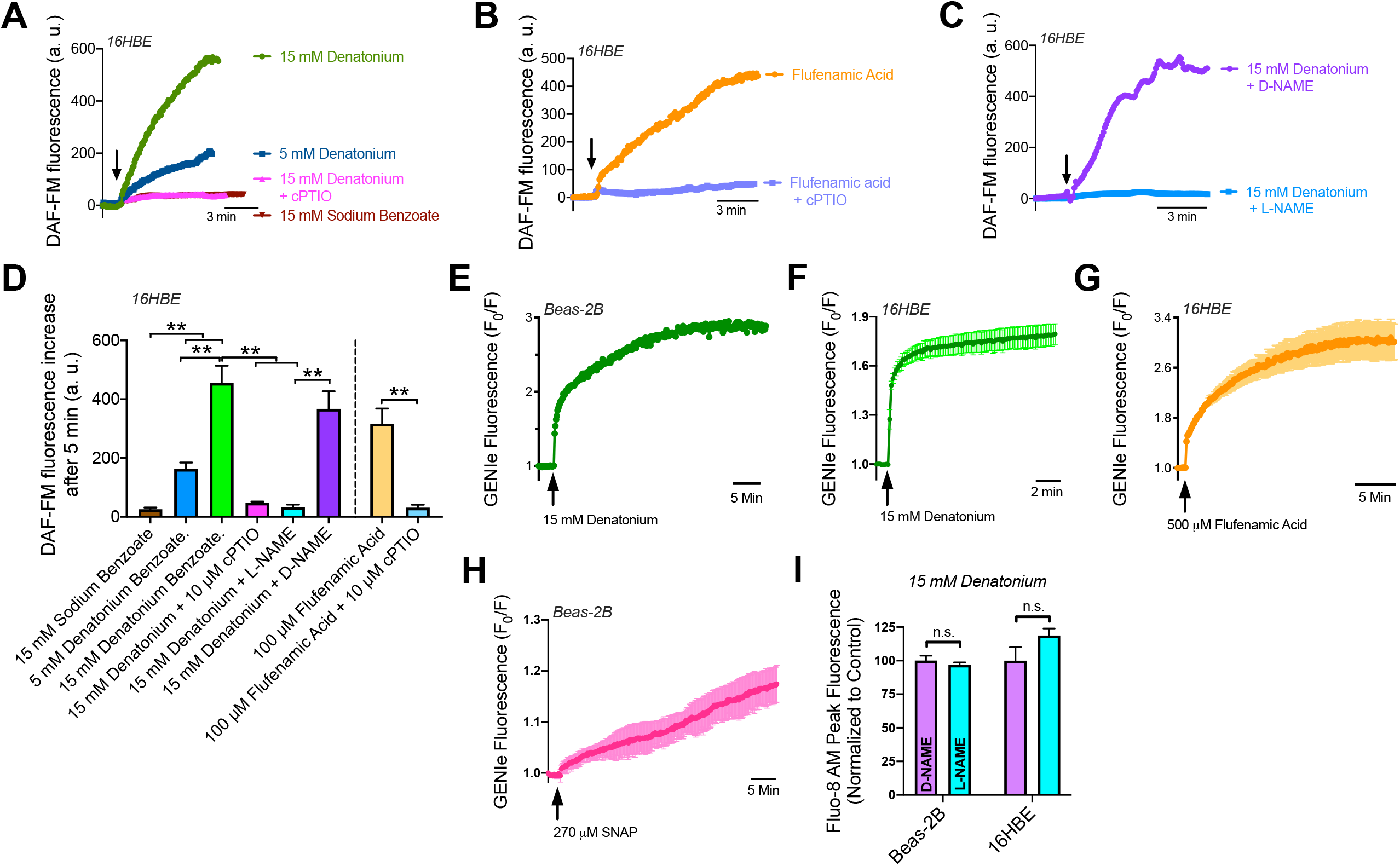
Non-ciliated airway cells produce nitric oxide in response to bitter compounds. **A-B** 16HBE cells were incubated with NO-sensing fluorescent probe, DAF-FM, which terminally reacts with NO and becomes fluorescent. Denatonium benzoate and FFA, but not sodium benzoate activated NO production that was blocked. By NO scavenger cPTIO. **C** NOS inhibitor, L-NAME, blocks denatonium-induced NO production. Negative control D-NAME had no effect. **D** Bar graph summarizing NO production after 5 min from 16HBE experiments as in *A-C*). **E-F** Cyclic guanosine monophosphate (cGMP) is produced when NO activates soluble guanylyl cyclase. Airway cells expressing cGMP biosensor, GENIe, increase cGMP, a downstream member of the NO pathway, in response to 500 μM flufenamic acid, 15 mM denatonium benzoate, or positive control 270 μM SNAP (NO donor). **I** NO production does not alter denatonium-induced Ca^2+^_i_ signaling in Beas-2B or 16HBE cells pretreated with L-NAME (100 μM, 1hr). Traces are representative experiments from 3 experiments. Data points on bar graph represent mean ± SEM from 3 experiments; significance determined by 1-way ANOVA using Sidak’s posttest.

### T2R14 and T2R39 localize partially to the nucleus

Intrigued by strong Ca^2+^_nuc_ responses and the partial nuclear staining of T2Rs above, we examined T2R localization in airway cells. GPCRs and associated proteins can localize to and function on the outer nuclear membrane or within internal nucleoplasmic reticulum membranes [32, 33]. A previous study reported an altered staining of T2R38 from cilia in normal tissue to nuclear localization in inflamed and de-ciliated CRS tissue [34], though antibody specificity was not verified nor were functional consequences reported. We hypothesized that certain T2Rs may localize to the nucleus or surrounding ER in the absence of proper trafficking to cilia.

Poor trafficking of T2Rs to the cell surface was reported in the context of potential requirement for β2 adrenergic receptors (β2ARs) to “chaperone” T2Rs [19]. However, most airway epithelial cells have robust β2AR expression; cAMP imaging with β2AR agonists in Beas-2Bs, 16HBEs, NCI-H292s, and A549s (**Supplementary Fig. S13**) suggests results here are not due to lack of β2ARs.

Immunofluorescence in multiple cells showed nuclear staining for T2R14 (responds to FFA, DPD, and quinine) and T2R39 (responds to denatonium benzoate and quinine) (**Fig. 4A-F, Supplementary Fig. S14**). The antibody against T2R14 was previously validated [6]. To test if the T2R39 antibody was specific, CRISPR-Cas9 was used to create a frameshift in TAS2R39 in A549s. No T2R39 staining was observed in these knockout cells, supporting nuclear localization (**Fig. 4B**). We also saw nuclear staining with an antibody against α-gustducin (**Figure 4A**), though implications are unclear in light of **Fig. 2E** and previous studies suggesting epithelial T2R signaling is gustducin-independent [35].

**Fig. 4.**
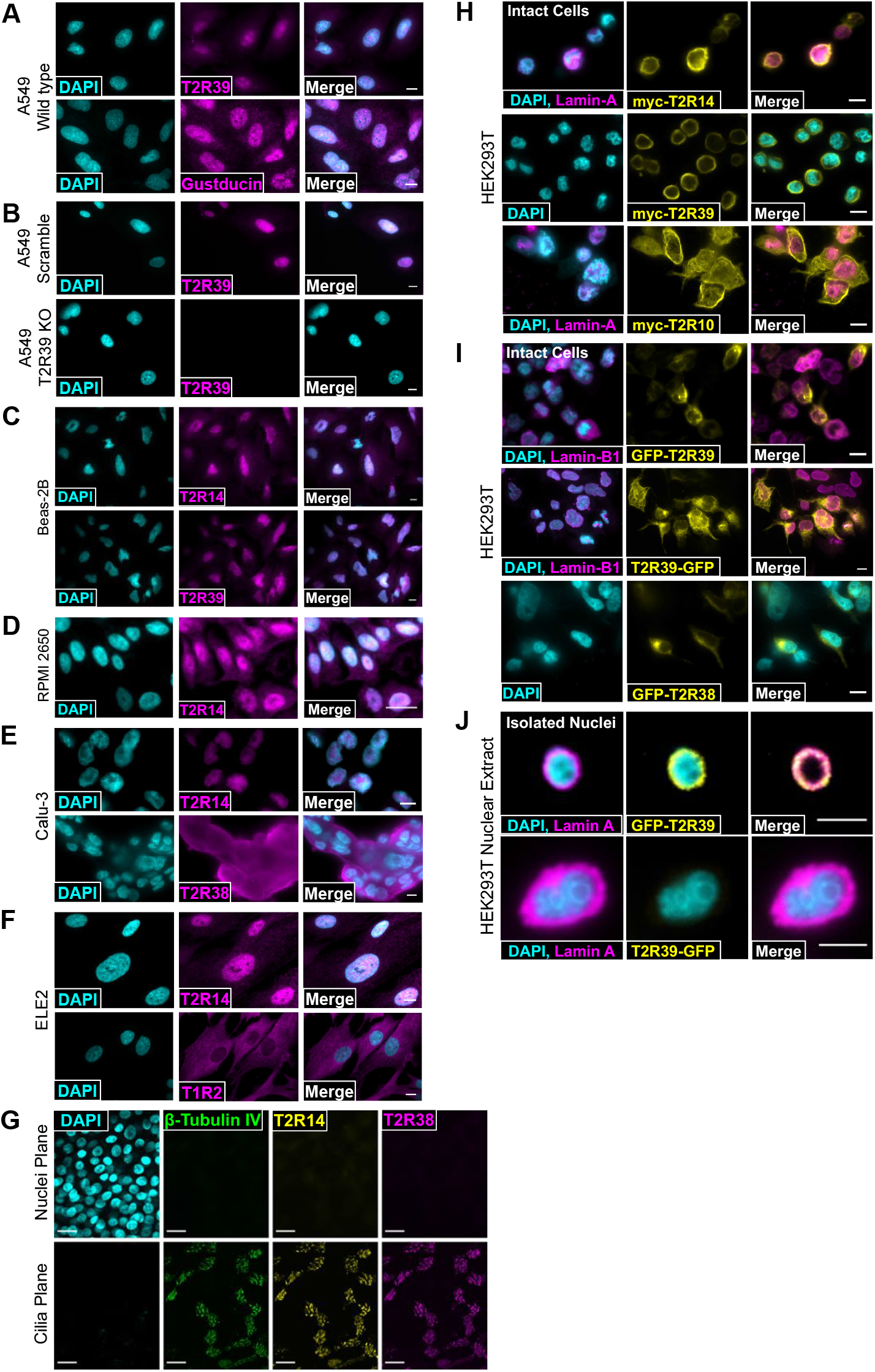
Nuclear localization of T2R14 and T2R39. **A-D** Fixed cultures of non-ciliated airway cells stained with antibodies targeting T2R14, T2R39, and Gustducin. CRISPR-cas9-induced scramble or T2R39 KO cell lines were stained for T2R39. **E-F** T2R38 and T1R2 are not localized to the nucleus in Calu-3 (lung adenocarcinoma) or ELE2 (hBMI immortalized primary bronchial epithelial cells) respectively. A-F scale bar is 10 µm. **G** Differentiated primary nasal epithelial cells contain T2R14 and T2R38 on cilia, not nuclei. All images are representative image from ≥3 independent experiments. Scale bar is 25 µm. **H** Fixed HEK cells expressing ectopic myc-tagged T2R14, T2R39, or T2R10 and co-expressing mCherry Lamin-A stained with anti-myc antibody, showing nuclear localization of T2R14 and T2R39. **I** Representative images showing GFP-T2R39 but not T2R39-GFP localizes to the nucleus in fixed HEK 239T cells. **J** Nuclei from HEK cells expressing GFP-T2R39 or T2R39-GFP. For all images, 1 representative image from ≥3 experiments were shown. For all images scale bars represent 10 µm.

In contrast, other GPCRs exhibited plasma membrane or cytoplasmic localization while T2R14 was nuclear (**Fig. 4E-F**). Denatonium-responsive T2R4, 8, and 39 are located at least partly to the nucleus via Western (**Supplementary Fig. S15**). The same T2R14 antibody showed T2R14 localization to cilia in differentiated primary nasal epithelial cells (**Fig. 4G**), as described [6]. T2R39 is likewise expressed in differentiated bronchial cilia [7]. Thus, in airway cells without cilia, such as de-differentiated squamous or cancer cells, T2R14 and T2R39 may instead localize at least partly to the nucleus.

Many studies utilize T2R expression constructs containing N-terminal rat somatostatin type 3 receptor (SSTR3) or bovine rhodopsin sequences to enhance plasma membrane localization (e.g., [19, 26]). While these constructs are useful for discovering T2R agonists, we tested localization of expressed T2Rs with minimal tagging (single N-terminal myc). We observed nuclear localization of myc-T2R14 and myc-T2R39, but not myc-T2R10 (**Fig. 4H**). Coupled with the endogenous differential T2R38 vs T2R14 localization (**Fig. 4E**), there may be different localizations of T2Rs within the same cells and between different cells (**Supplementary Fig. S15**).

We also expressed either N-terminal or C-terminal GFP fusions. N-terminally tagged GFP-T2R39 co-localized partly with nuclear membrane Lamin-B1, while C-terminally tagged T2R39-GFP appeared less nuclear. C-terminal sequences in mGluR5 are important for nuclear localization [36]. Similar C-terminal sequences may be important for T2R39 and C-terminal GFP may block interactions conferring nuclear localization. GFP-T2R38 did not appear nuclear, suggesting that the nuclear localization of GFP-T2R39 is an effect of the T2R39 sequence and not the N-terminal GFP (**Fig. 4I**). To further test this, we co-expressed either GFP-T2R39 or T2R39-GFP with mCherry-Lamin A in HEK293Ts. Consistent with above, GFP-T2R39 was localized to the membrane of isolated HEK293T nuclei (labeled with mCherry-Lamin A) while T2R39-GFP was not (**Fig. 4J**), supporting a role for C-terminal sequences in T2R39 nuclear localization.

### Ca^2+^_nuc_ signals to mitochondria

What are the consequences of T2R-induced Ca^2+^_nuc_ in non-ciliated airway cells? In airway smooth muscle cells, T2R agonists induce phosphorylation of p38 MAPK [37], important for both cell survival. Western revealed that DPD, FFA, and denatonium benzoate promoted a 10-25-fold increase in p38 phosphorylation (**Supplementary Fig. S16A-C**). However, buffering Ca^2+^ with BAPTA did not block p38 MAPK phosphorylation (**Supplementary Fig. S16D**), suggesting that Ca^2+^_i_ and p38 MAPK activation are independent.

Mitochondria are in close proximity to nuclei (**Fig. 5A**). Bitterant absinthin activates influx of mitochondrial Ca^2+^ (Ca^2+^_mito_) [38]. We observed sustained, mitochondrial-reminiscent Ca^2+^ elevations in submerged primary nasal cells (**Supplementary Fig. S7**). Thus, we hypothesized that elevation of Ca^2+^_nuc_ might signal to mitochondria. We expressed mitochondrial-localized Ca^2+^ sensor 4mtD3cpv [39] in 16HBEs, Beas-2Bs, or HEKs. 16HBEs treated with thapsigargin in 0-Ca^2+^_o_ buffer exhibited increased Ca^2+^_mito_, suggesting internal Ca^2+^ stores can feed mitochondria (**Fig. 5B**). Denatonium benzoate and FFA also increased Ca^2+^_mito_ in Beas-2Bs (**Fig. 5C-D**) and 16HBEs (**Fig. 5F**) but not HEK293Ts (**Fig. 5E**), which do not exhibit Ca^2+^_i_ increases to denatonium. This was blocked by U73122 or IP_3_ receptor antagonist Xestospongin C (**Fig. 5G-H**). Thus, bitterants activate Ca^2+^_nuc_ and Ca^2+^_mito_ with similar signaling requirements. **Ca**^**2+**^_**nuc**_ **activates cell death**

**Fig. 5.**
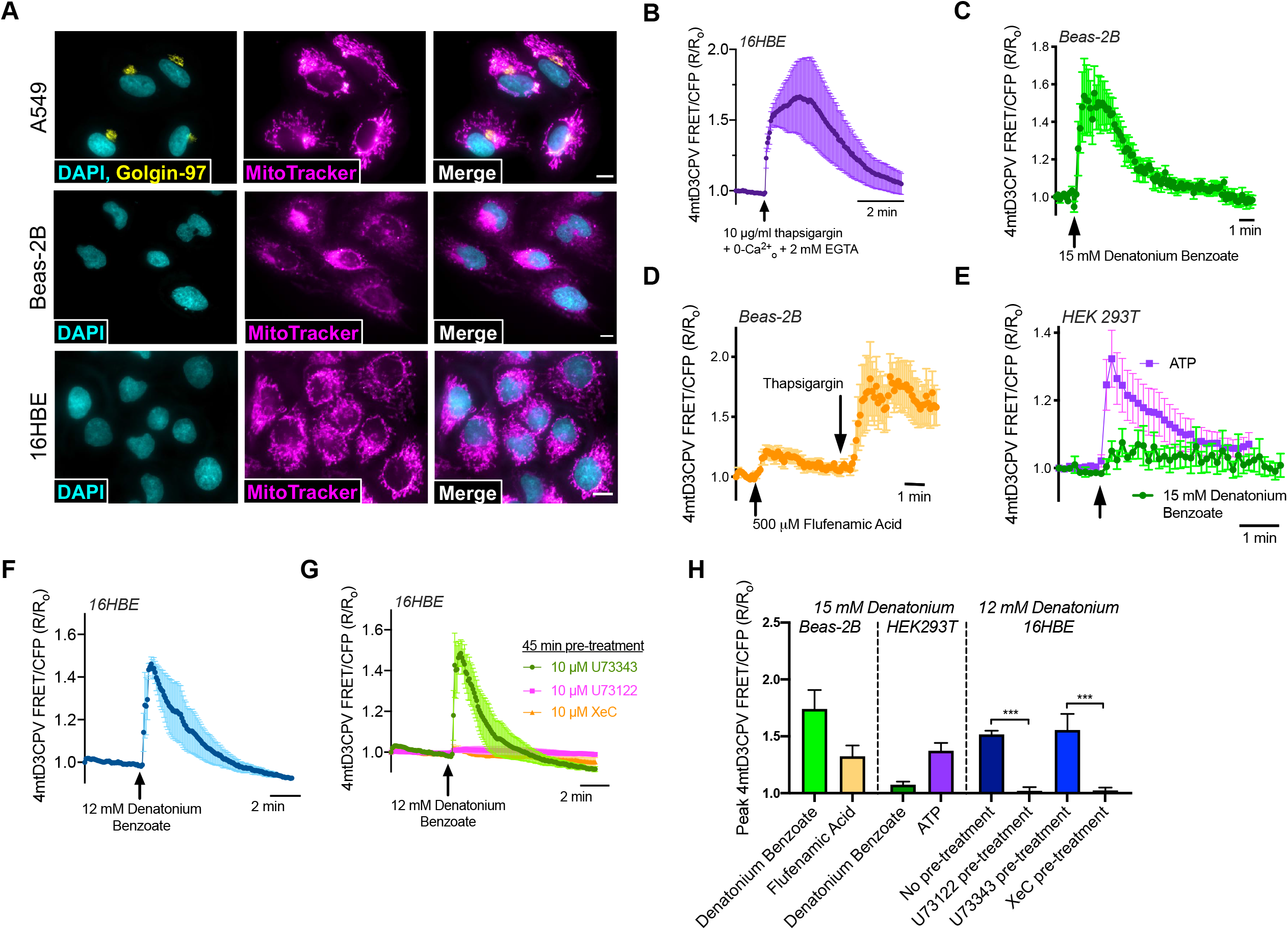
Acute transient Ca^2+^_mito_ elevation in airway cells treated with T2R agonists. **A** Fluorescence images of nuclear stain DAPI and MitoTracker in A549, Beas-2B, and 16HBE cells, plus immunofluorescence for Golgi marker golgin-97 in A549 cells. Scale bars are 10 µm. **B** Trace of ratiometric CFP/YFP FRET-based mitochondrial-targeted Ca^2+^ indicator 4mtD3cpV in 16HBEs showing elevation of intracellular Ca^2+^ by Ca^2+^ ATPase inhibitor thapsigargin in 0-Ca^2+^_o_ HBSS (no added Ca^2+^ and 2 mM EGTA) increased mitochondrial Ca^2+^, demonstrating that store Ca^2+^ release can elevate Ca^2+^_mito_. **C**,**D** Trace showing stimulation of Beas-2Bs with denatonium or flufenamic acid increased Ca^2+^_mito_. **E** Trace showing minimal Ca^2+^_mito_ response in HEK cells, which also do not exhibit Ca^2+^_nuc_ responses to denatonium. **F** Trace showing denatonium elevated Ca^2+^_mito_ in 16HBE cells. **G** This elevation of Ca^2+^_mito_ was blocked by pre-treatment with PLC inhibitor U72122 or IP_3_R inhibitor xestospongin C (XeC). Note that DMSO concentration (0.1%) is the same for all conditions, so inactive U73343 pretreatment also serves as vehicle control for XeC. **H** Bar graph showing peak 4mtD3CPV responses represented as mean ± SEM from independent experiments shown in **B-G**. All traces show representative results from ≥4 transfected cells from one of ≥4 independent experiments imaged at 60x.

Overload of Ca^2+^_mito_ is linked to cell death [40]. Denatonium benzoate, but not sodium benzoate, inhibited XTT reduction (indirect measurement of NADH metabolism; **Fig. 6A**) at concentrations ≥1 mM in A549’s (**Fig. 6B,C**). T2R agonist chrysin also impaired XTT reduction (**Fig. 6C**). In 16HBEs and Beas-2Bs, denatonium benzoate and quinine, but not sodium benzoate, reduced mitochondrial potential (ΔΨ_m_), measured by JC-1 (**Fig. 6D-G**). T2R agonists quinine, 3-oxo-C12-HSL, and DPD activated apoptosis, visualized by increased fluorescence of caspase 3/7-sensitive DEVD-based dye CellEvent (**Fig. 6H and I**). Apotosis-activating concentrations of denatonium (>5 mM; **Fig. 6H**) paralleled concentrations eliciting Ca^2+^_nuc_ (**Fig. 6J**) and altering ΔΨ_m_ (**Fig. 6F**). Pre-incubation of Beas-2Bs in 0-Ca^2+^ HBSS with 10 μM BAPTA ± 15 mM denatonium prevented both ΔΨ_m_ depolarization (**Fig. 6K**) and caspase activation (**Fig. 6L**) over 6 hours. Thus, bitterant-induced Ca^2+^ elevation is required for ΔΨ_m_ depolarization and apoptosis.

**Fig. 6.**
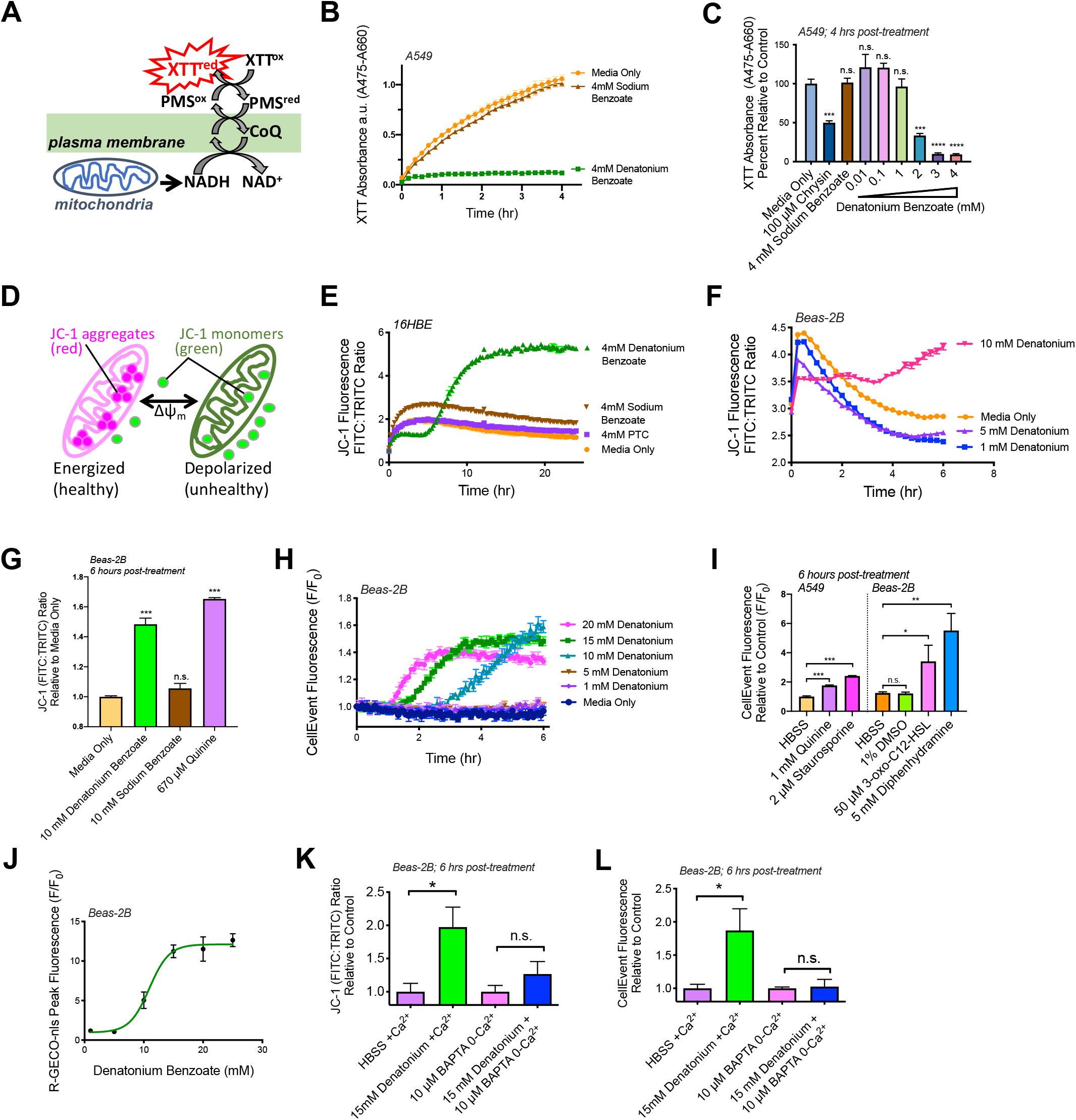
Denatonium halts cell proliferation, depolarizes mitochondrial membrane potential (Ψ_m_), and initiates apoptosis. **A** Depiction of XTT assay. **B** Denatonium, not benzoate, halts A549 metabolism. **C** A549 metabolism is impaired by T2R agonists chrysin and denatonium at concentrations >1 mM. **D** Mitochondrial membrane potential detection dye JC-1 shifts from red to green fluorescence with depolarization. **E-G** Traces and bar graph of ratiometric mitochondrial membrane potential dye (JC-1) showing that denatonium, not benzoate, initiates mitochondrial membrane depolarization in 16HBEs (*E*) and Beas-2Bs (*F-G*). **H** Trace showing fluorescence increases signifying caspase activation in Beas-2Bs were incubated with denatonium and CellEvent caspase-3/7 detection. **I** Bar graph CellEvent fluorescence at 6 hours, indicating caspase activation in A549s in response to quinine and in Beas-2Bs in response to 3-oxo-C12HSL and diphenhydramine. Staurosporine = positive control. **J** Does response of peak Ca^2+^_nuc_ (R-GECO-nls) in Beas-2B treated with denatonium. **K-L** Beas-2Bs were pre-incubated with 10 μM BAPTA in 0-Ca^2+^ HBSS then treated with denatonium in JC-1 (*K*) or CellEvent (*L*) assays. Time course experiments are representative of ≥3 independent experiments. Bar graphs show mean ± SEM from ≥3 experiments. Significance by 1-way ANOVA using Dunnett’s or Tukey’s posttest \**P*<0.05, \*\*\**P*<0.0005 \*\*\*\**P*<0.0001.

To test physiological relevance, we examined bitterant-induced apoptosis in de-ciliated squamous primary human bronchial epithelial (HBE) cells exposed to apical submersion and fully differentiated ciliated cultures exposed to apical air (**Fig. 7A and Supplementary Fig. S17**). Bitter agonists increased Annexin V staining over 3-6 hrs in squamous but not ciliated cultures (**Fig. 7B-C**). To tie these responses to T2Rs, similar differentiated vs squamous primary nasal cultures were stimulated with T2R38 agonist PTC. PTC increased annexin V staining only in squamous cultures homozygous for the functional T2R38 (PAV polymorphism). Cultures homozygous for non-functional T2R38 (AVI polymorphism) did not exhibit increased annexin V staining (**Fig. 7D-E**), suggesting increased Annexin V-staining is dependent on functional T2R38. Submerged squamous HBEs also showed staining with propodium iodide (PI; reflecting permeabilization) 6 hours after bitterant stimulation (**Fig. 7F-G**), likely reflecting secondary necrosis in the absence of phagocytes [41].. The earlier onset of Annexin V versus PI staining supports apoptosis. These data suggest that T2R agonists activate apoptosis in squamous but not well-differentiated epithelial cells.

**Fig. 7.**
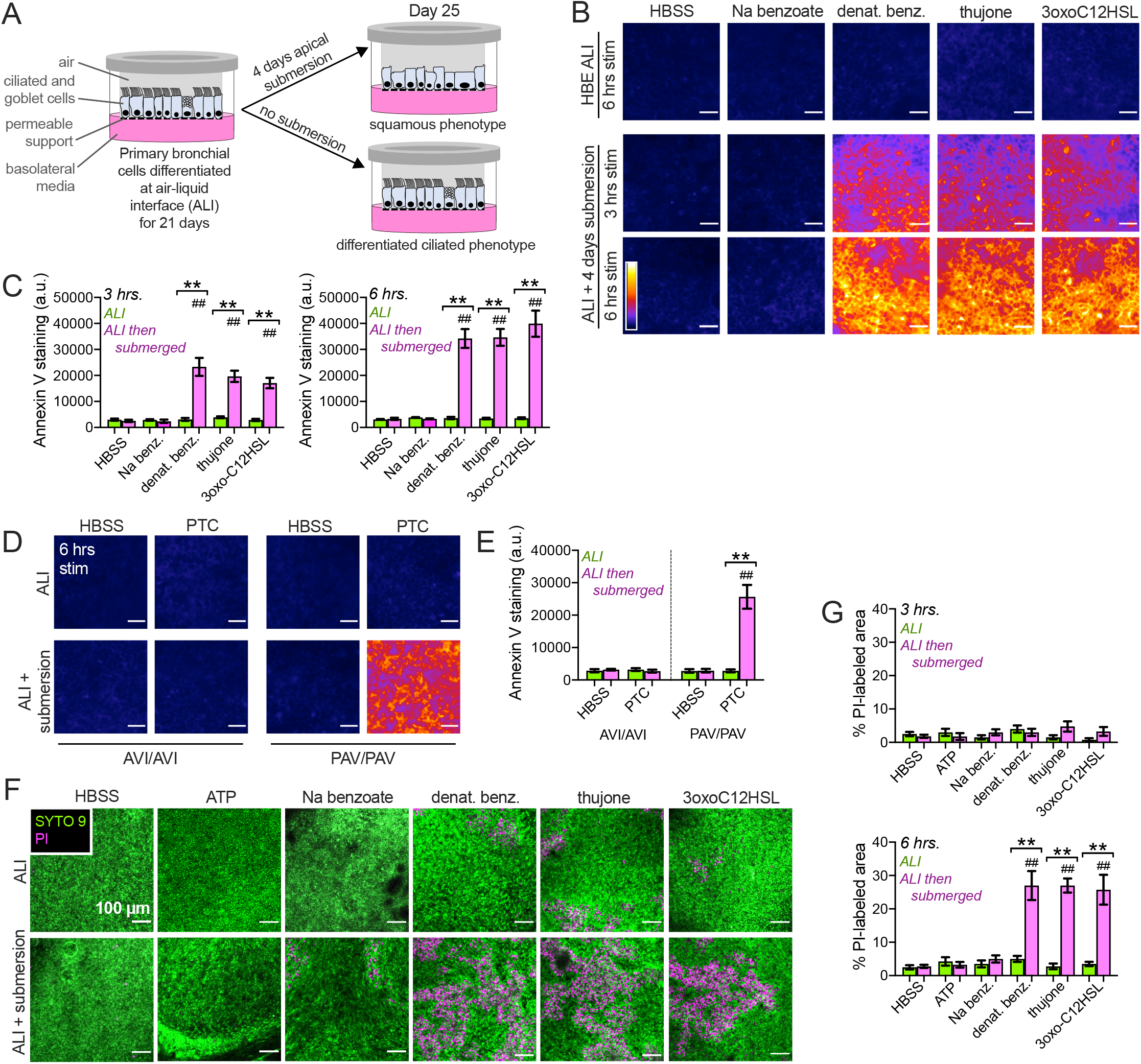
T2R agonists induce cell death in submerged but not differentiated primary HBE and sinonasal cells. **A** Primary HBE cells grown on transwells and differentiated at air liquid interface (ALI) were exposed to 4 days of air or apical submersion (as described in the Methods and [17]). Apical submersion increases squamous differentiation ([17, 70, 71] and **Supplementary Fig. S17**). **B** Denatonium benzoate, thujone, or 3-oxo-C12HSL increased Annexin V FITC staining in deciliated (submerged) but not differentiated HBE cells. Representative intensity pseudocolored images shown. Scale bar 100 µm. Sodium benzoate (control for denatonium benzoate) had no effect. **C** Bar graphs of relative fluorescence intensity (mean ± SEM) from 4 cultures (2 each from 2 donors) per condition stained at 3 (left) or 6 hours (right). Green bars show air-exposed; magenta bars show submerged. Significance by 1-way ANOVA, Bonferroni posttest; \*\**p*<0.01 between bracketed bars (submerged vs air-exposed) and ^##^*p*<0.01 vs HBSS only. **D** Experiments carried out similarly to *A-B* in *TAS2R38*-genotyped primary nasal cultures ± 500 µM PTC. *TAS2R38* (encoding T2R38) has two polymorphisms Mendelianly-distributed in the Philadelphia population [72]. The PAV allele encodes a functional receptor while the AVI allele encodes a non-functional receptor [73]. ALIs from PAV/PAV patient cells exhibit Ca^2+^ responses to T2R38-specific agonist PTC while ALIs from AVI/AVI homozygous patient cells do not [3]. Representative images show Annexin V-FITC staining at 6 hrs, which increased in PAV/PAV cells exposed to submersion but not air-exposed cells. Staining did not increase in AVI/AVI cells under either condition. **E** Bar graph of mean ± SEM of Annexin V FITC staining from 4 cultures per condition, each from a separate PAV/PAV or AVI/AVI patient. Significance by 1-way ANOVA with Bonferroni posttest; \*\**p*<0.01 between bracketed bars (submerged vs air-exposed) and ^##^*p*<0.01 vs HBSS only. **F** Representative images of live-dead (Syto9 in green and propidium iodide [PI] in magenta) staining of HBE cells. **G** Bar graph (mean ± SEM,4 cultures per condition, 2 each from 2 donors) of PI-labeled area from experiments as in *E*. Significance by 1-way ANOVA, Bonferroni posttest; \*\**p*<0.01 between bracketed bars (submerged vs air-exposed) and ^##^*p*<0.01 vs HBSS only.

## Discussion

Some T2Rs are localized to cilia of differentiated cells [3, 7] but may localize to the nucleus in less differentiated cells. A prior study reported nuclear localization of T2R38 in tissue from CRS patients [34], though no function was reported. We show that T2R14 and T2R39, likely located on the nucleus of non-ciliated airway cells, signal through Ca^2+^_nuc_, cAMP_nuc_, and NO to increase Ca^2+^_mito_ and depolarize ΔΨ_m_, initiating apoptosis. Bitterants [42-45] and bacterial products [46, 47] activate apoptosis and denatonium alters ΔΨ_m_ [45], but was attributed to oxidative stress [45]. We suggest that T2R activation elevates Ca^2+^_nuc_ and Ca^2+^_mito_ to activate ΔΨ_m_ depolarization to initiate apoptosis. We hypothesize that intense Ca^2+^_nuc_ increase in close proximity to mitochondria overloads Ca^2+^_mito_.

We hypothesize that absence or loss of motile cilia causes some T2Rs to alternatively localize to nuclei. Both endogenous or heterologous T2R14 and T2R39 can localize to the nucleus. Many GPCRs localize to the nucleus and regulate processes like proliferation or synaptic function [32, 48]. Some nuclear GPCRs may act as transcription factors [49]. Nuclear GPCRs may have unique localization sequences [48]; mGluR5 receptor localizes to the nucleus through C-terminal sequences [36]. The ratio of plasma membrane-to-nuclear mGluR5 varies among neuronal types [50], suggesting that trafficking depends on additional factors. Masking the C-terminus of T2R14 or T2R39 with GFP reduced nuclear localization, suggesting C-terminal interactions are important. Future studies must identify sequences important for nuclear and/or cilia localization. Transport of proteins into cilia and nuclei involves shared components [51], including importins and nucleoporins [52-54]. This may be due to movement of signaling molecules from the cilia to the nucleus during primary ciliary signaling and regulation of transcription [55]. We hypothesize that, in de-ciliated airway cells, the nuclear membrane may become a reservoir for GPCRs that would normally traffic to the cilia under normal conditions.

Non-ciliated squamous airway cell T2Rs retain the ability to activate NO production, as observed during activation of T2Rs in ciliated cells [3, 5]. While eNOS is localized to the cilia base in airway epithelial cells [56, 57], eNOS localizes to the nucleus in some cells [58, 59]. Altered localization of eNOS to the nucleus by loss of cilia may facilitate coupling T2Rs to NO despite altered T2R localization.

Many bitterants are hydrophobic, allowing them to activate intracellular T2Rs. Quinine rapidly permeates taste and other cells [60, 61]. Bacterial agonists like 3-oxo-C12-HSL are cell permeant [62] and reach extracellular concentrations up to hundreds of µM in late stationary cultures of *P. aeruginosa* [63]. Colonization of bacteria during airway infection could generate localized AHL levels high enough to enter the cell and activate T2R Ca^2+^_nuc_. Ca^2+^_nuc_ is linked to proliferation [64] and regulation of transcription [65]. Mitochondrial-to-nuclear Ca^2+^ signaling may regulate gene expression [66, 67]. We show that reverse nuclear-to-mitochondrial Ca^2+^ can regulate apoptosis.

Apoptosis is part of innate immune defense [68] and may be a “last resort” response to bacterial metabolites during sustained infection or during diseases involving epithelial remodeling, but may also allow bacteria to damage inflamed or remodeled epithelia as bitter bacterial metabolites activate T2Rs, Ca^2+^_nuc_, and apoptosis in de-ciliated cells. There are also potentially important implications for epithelial-derived lung cancers. We report Ca^2+^_nuc_ and apoptosis in lung cancer lines but not in well-differentiated primary cells. T2R agonists may therapeutically activate apoptosis in tumor but not differentiated cells. T2R activation by bacterial metabolites may also contribute to tumor-microbiome crosstalk. Whether activation of T2R-induced Ca^2+^_nuc_ and apoptosis is detrimental or beneficial in non-ciliated epithelial cells thus likely depends on disease context.

## Materials Availability

All materials will be provided upon reasonable request to RJL.

## Acknowledgements

We thank Maureen Victoria (University of Pennsylvania) for technical assistance and discussion and Andrew Ramsey (University of Pennsylvania) and Alfred Sloan (Synthego) for helpful discussion.

## Author Contributions

Conceptualization and Visualization: D.B.M. and R.J.L.; Investigation and Formal Analysis: D.B.M., L.E.K., M.E.J., P.O.J., R.L.J.H., R.M.C., and R.J.L.; Writing – Original Draft: D.B.M.; Writing – Review and Editing: D.B.M. and R.J.L.; Data Curation and Resources: N.D.A. and J.N.P.; Funding Acquisition and Supervision: R.J.L.

