## Supplementary Figures for "The bitter end: T2R bitter receptor agonists elevate nuclear calcium and induce apoptosis in non-ciliated airway epithelial cells"

**Detailed Materials and Methods**

***Human primary cell isolation and culture***

Primary human sinonasal cell culture was carried out as extensively described [1, 2, 3]. Tissue acquisition was done in accordance with The University of Pennsylvania guidelines for the use of residual clinical material and in accordance with the U.S. Department of Health and Human Services code of federal regulation Title 45 CFR 46.116 and the Declaration of Helsinki. Institutional review board approval (#800614) and written informed consent from each patient was obtained. Tissue was used from patients ≥18 years of age undergoing surgery for sinonasal disease (CRS) or other procedures (e.g. trans-nasal approaches to the skull base for pituitary tumors).

Human primary nasal epithelial cells were obtained through enzymatic dissociation of human sinonasal tissue and cultured as described [4]. Human primary bronchial epithelial cells were procured commercially (Cat.# CC-2540S, Lonza). For isolation of primary nasal epithelial cells, patient sinonasal specimens were digested in MEM culture medium containing 1.4 mg/ml protease and 0.1 mg/ml DNase for 1 hour at 37°C followed by 2 neutralizing washes with MEM culture medium containing 10% fetal bovine serum. The cell suspension was transferred to a T-25 culture flask containing PneumaCult-Ex Plus culture medium (Cat.# 05040, Stemcell Technologies) supplemented with 100 U/ml penicillin and 100 μg/ml streptomycin and incubated at 37°C, 5% $CO_2$ for 2 hours to allow for adherence and removal of non-epithelial cells (e.g. fibroblasts, macrophages and lymphocytes). The cell suspension containing only nasal epithelial cells was then transferred to a 10 cm tissue culture dish containing PneumaCult-Ex Plus culture medium and left in the incubator overnight. The following day, the culture medium was aspirated and the cells washed once with sterile PBS and replaced with fresh PneumaCult-Ex Plus culture medium. Both nasal and bronchial primary epithelial cells were passaged every 4-5 days or when cultures achieved 80% confluence. Primary human bronchial epithelial cells were obtained from Lonza and cultured as above.

For deciliation experiments, Normal human bronchial epithelial cells (HBE) air liquid interface cultures (ALIs) were differentiated for 3 weeks and either exposed to air or subjected to submersion for 4 days as previously described [2]. All comparisons were made between age-matched cultures. Cells were stimulated apically for 3-6 hours with 10 mM sodium benzoate (Na benzoate), 10 mM denatonium benzoate (denat. benz.), 600 µM thujone, or 100 µM 3oxoC12HSL or no stimulation HBSS alone. Cells were then stained with annexin V-FITC assay kit (Cayman Chemical Cat. # 600300). Fluorescence was immediately imaged at the center of the ALI on a wide-field fluorescence microscope with 10x objective. TAS2R38 genotype was carried out as previously described [4]. The *TAS2R38* gene encoding the T2R38 receptor has two common polymorphisms that are Mendelianly-distributed in the Philadelphia population [5, 6, 7]. The PAV allele encodes a functional receptor while the AVI allele encodes a non-functional receptor [8]. ALIs from PAV/PAV homozygous patient cells exhibit $Ca^{2+}$ responses to T2R38-specific agonist PTC while ALIs from AVI/AVI homozygous patient cells do not [4].

#### *Cell line culture*

Cell line culture was as described [1, 2, 3]. RPMI 2650, BEAS-2B, Calu-3, A549, HEK293T, NCI-H292, and Caco-2 cells were obtained from ATCC (Manassas, VA USA). 16HBE (SV-40 immortalized normal human bronchial) cells [9] were obtained from D. Gruenert (University of California San Francisco, San Francisco, CA USA). Cells were grown in submersion in Minimal Essential Media with Earl's salts (Gibco; Gaithersburg, MD USA) supplemented with 10% FBS, 1% penicillin/streptomycin mix (Gibco). Transfection of submerged BEAS-2B and A549 cells was carried out in 8-well chamber slides (CellVis) with Lipofectamine 3000 as per the manufacturer's instructions.

#### *Live cell imaging*

For $Ca^{2+}$ imaging, cells were loaded with either 5 µM of either Fura-2-AM or Fluo-8-AM for 1 hour in HEPES-buffered Hank's Balanced Salt Solution at room temperature in the dark. Fura-2 was imaged using an Olympus IX-83 microscope (20x 0.75 NA PlanApo objective), fluorescence xenon lamp (Sutter Lambda LS, Sutter Instruments, Novato, CA USA), and excitation and emission filter wheels (Sutter Lambda 2). Images were acquired with an Orca Flash 4.0 sCMOS camera (Hamamatsu, Tokyo, Japan). Images were acquired with MetaFluor (Molecular Devices, Sunnyvale, CA USA) using a standard Fura-2 filter set for dual excitation (79002-ET, Chroma Technologies, Rockingham, VT USA). Imaging of Fluo-8 was performed similarly to Fura-2 except excitation utilized 470/40 nm excitation filter, 495 lp dichroic, and 525/40 nm em filter (49002-ET, Chroma Technologies) with a XCite 120 LED Boost excitation light source (Excelitas Technologies). For NO, cells were loaded with 10 uM DAF-FM diacetate for 1.5 hours ± 5 µM cPITO then incubated ±L- or D-NAME prior to imaging. DAF-FM was imaged identically to Fluo-8. For nuclear $Ca^{2+}$ or cAMP either $Ca^{2+}$ biosensors GECO or

R-GECO-nls [10], or cAMP biosensors Flamindo2 or nls-Flamindo2 [11] were transfected using lipofectamine 3000 (ThermoFisher Scientific) 24-72 hours prior to imaging. Live cell images were taken on an Olympus IX-83 microscope using x20 objective (0.8 NA; MetaFluor software) with standard FITC or TRITC filter set and microscope set up as above.

***Proliferation, Mitochondrial Membrane Potential, and Apoptosis Measurements***

Cells were cultured at sub-confluent density in 24-well glass bottom plates (CellVis). XTT (sodium 3´-[1- (phenylaminocarbonyl)- 3,4- tetrazolium]-bis (4-methoxy6-nitro) benzene sulfonic acid hydrate) was added immediately before absorbance measurements at 475nm (specific absorbance) and 660nm (reference absorbance). JC-1 dye (etraethylbenzimidazolylcarbocyanine iodide) was added 10 min prior to measurements (ex.488/em.535 and em.590) while CellEvent Caspase 3/7 was added directly prior to measurements (ex.495/em.540) as per manufacturer's specifications. All data were obtained using a Spark 10M fluorescence plate reader (Tecan; Männedorf, Switzerland).

***Immunofluorescence microscopy***

Cultures were fixed in 4% paraformaldehyde for 20 min at room temperature followed by a simultaneous blocking/permeabilization in phosphate saline buffer containing 5% normal donkey serum, 1% bovine serum albumun, 0.2% saponin, and 0.1% Triton X-100 for 45 min at room temperature. Cultures were incubated in primary antibody, 1:100 dilutions of T2R or α-gustducin antibodies, at 4°C overnight. Cultures were then incubated with AlexaFluor-labeled donkey anti-mouse or anti-rabbit (1:1000) at 4°C for 1 hour then mounted with Fluoroshield with DAPI (Abcam). All microscopy images were taken on an Olympus IX-83 microscope using x60 objective (1.4 NA oil; MetaMorph software). For ectopic expression: Myc-T2R10, myc-T2R39, GFP-T2R39, and T2R39-GFP expression constructs obtained from VectorBuilder (Chicago, IL). Cells were transfected using Lipofectamine 3000. Thermo MitoTracker Deep Red FM used at 10 nM for 15m pre-incubation at 37° C before fixing with 4% PFA then staining DAPI ± Golgin-97 antibody (Life Technologies, A21270, 1:100; anti-mouse AF488 secondary 1:1000). For co-staining of T2R14 and T2R38 in primary cell cilia, rabbit primary antibodies were labeled with Xenon AlexaFluor 488 or 546 conjugated fab fragments using Zenon antibody labeling kit (ThermoFisher; Cat # Z25302 and Z25304, respectively).

***Generation of Stable shRNA Cell Lines***

Beas-2Bs were transfected with pRS plasmids (OriGene, Rockville, MD) to express shRNA for either T2R8 (T301219B), T2R10 (T301240B), T2R14 (TR301238A and TR301238B), or scramble shRNA (TR30012). Twenty-four hours after transfection, cells were supplemented with media containing 1 µg/mL puromycin for 1 week, and then maintained with media containing 0.2 µg/mL puromycin prior to experiments.

***Biochemistry and Western Blotting***

For endogenous T2R Western blot analysis, cells were lysed in 50 mM Tris pH 7.5, 150 mM NaCl, 1% IGEPAL CA-630, 1% deoxycholate, 1 mM NaF, 1 mM DTT, DNase, and Protease Inhibitor Cocktail (Roche cOmplete). Post 800 x *g* lysates (60 *µ*g protein/lane) were loaded in a NuPage 4-12% Bis-Tris gel. Separated proteins were then transferred to nitrocellulose. Membrane was blocked in 5% milk in 50 mM Tris, 150 mM NaCl, and 0.025% Tween-20 (Tris-Tween) for 1 hour. Primary antibody was used at 1:1000 dilution in Tris-Tween with 5% BSA for 1.5 hours. Secondary antibodies of goat anti-rabbit or anti-mouse IgG-horseradish peroxidase were diluted 1:5000 in blocking buffer then incubated with membrane for 1 hour. Blots were incubated for 5 min with Clarity ECL (BioRad Laboratories) and images were obtained using a BioRad Gel Doc, subsequent images and densitometry was analyzed using Image Lab Software (BioRad Laboratories). Nuclei were isolated using the REAP method [12] then either used for biochemistry or fixed on glass slides (4% formaldehyde in PBS, 15 min), stained as above, and imaged.

***Quantitative PCR (qPCR)***

Subconfluent cultures were resuspended in TRIzol (ThermoFisher Scientific) and used immediately or otherwise stored at -70°C until use. RNA was isolated using Direct-zol RNA kit (Zymo Research). Once purified, RNA was transcribed to cDNA via High-Capacity cDNA Reverse Transcription Kit (ThermoFisher Scientific). Resulting cDNA was then quantified utilizing Taqman Q-PCR probes in a QuantStudio 5 Real-Time PCR System (ThermoFisher Scientific). Data was then analyzed using Microsoft Excel and plotted in GraphPad PRISM.

***Data analysis and statistics***

T-tests (two comparisons only) and one-way ANOVA (>2 comparisons) were calculated using GraphPad PRISM with appropriate post-tests. For comparisons of all samples, Tukey-Kramer post-test was used, while Bonferonni post-test was used for selected pairwise comparisons. Any other data analysis was performed in Microsoft Excel. In all figures, $p < 0.05$ (*), $p < 0.01$ (**), $p < 0.001$ (***), while no statistical significance was represented by “n.s.” All data points represent the mean ± SE

### Supplementary Table S1: Key biological and chemical reagents used in the study.

| Reagents | Source | Product Number |
|---|---|---|
| *Antibodies* | | |
| anti-Lamin A + Lamin C | Abcam | ab108595 |
| anti-T2R8 | Abcam | ab75109 |
| anti-T2R38 | Abcam | ab65509 |
| anti-b-tubulin IV | Abcam | ab11315 |
| anti-T2R38 | Abcam | ab130503 |
| anti-T1R2 | ThermoFisher Scientific | PA5-34254 |
| anti-T2R39 | ThermoFisher Scientific | PA5-39710 |
| GNAT3 | ThermoFisher Scientific | PA5-50670 |
| anti-T2R4 | ThermoFisher Scientific | PA-67752 |
| anti-T2R14 | ThermoFisher Scientific | PA5-39710 |
| anti-Lamin B2 clone E-3 | ThermoFisher Scientific | 332100 |
| AlexaFluor (AF) 488-conjugated donkey anti-mouse IgG | ThermoFisher Scientific | A21202 |
| AF546-conjugated donkey anti-mouse IgG | ThermoFisher Scientific | A10036 |
| AF546-conjugated donkey anti-rabbit IgG | ThermoFisher Scientific | A10040 |
| AF647-conjugated donkey anti-rabbit IgG | ThermoFisher Scientific | A31573 |
| anti-α-tubulin | Developmental Studies Hybridoma Bank | 12G10 |
| anti-Myc-tag | Cell Signaling Technology | 9B11 |
| anti-phospho-p38 MAPK | Cell Signaling Technology | 9211 |
| anti-Golgin 97 | Life Technologies, | A21270 |
| *Bacterial & Viral Strains* | | |
| Green downward cGMP sensor GENIe BACMAM | Montana Molecular | D0800G |
| T2R39 CRISPR knockout lentivirus (vector info below) | Cyagen Biosciences | N/A |
| Scramble CRISPR lentivirus (vector info below) | Cyagen Biosciences | N/A |
| hBMI Lentivirus (vector info below) | Cyagen Biosciences | N/A |
| *Chemical, Agonists, and Inhibitors* | | |
| U73122 | Cayman Chemical | 70740 |
| U73343 | Cayman Chemical | 17339 |
| L- and d-$N^G$-nitroarginine methyl ester (L-NAME) | Cayman Chemical | 80210 |
| D-$N^G$-nitroarginine methyl ester (D-NAME) | Cayman Chemical | 21687 |
| Apigenin | Cayman Chemical | 10010275 |
| Parthenolide | Cayman Chemical | 70080 |
| Diphenhydramine | Cayman Chemical | 11158 |
| Histamine | Millipore Sigma | H7125 |
| ATP | Millipore Sigma | A9187 |
| (-)-a-Thujone | Millipore Sigma | 89231 |
| Flufenamic acid (FFA) | Millipore Sigma | F9005 |
| Denatonium benzoate | Millipore Sigma | D5765 |

| | | |
|---|---|---|
| Phenylthiocarbamide (PTC) | Millipore Sigma | P7629 |
| Sodium Benzoate | Millipore Sigma | B3420 |
| Diphenidol | Millipore Sigma | SML2169 |
| Quinine | Millipore Sigma | Q0132 |
| 3-oxo-C12HSL | Millipore Sigma | O9139 |
| Saponin | Millipore Sigma | S7900 |
| Bovine Serum Albumin | Millipore Sigma | A2153 |
| b-escin | MP Biomedicals | 0215794101 |
| Fura-2-AM | ThermoFisher Scientific | F1221 |
| JC-1 | ThermoFisher Scientific | T3168 |
| DAF-FM diacetate | ThermoFisher Scientific | D23844 |
| BAPTA-AM | ThermoFisher Scientific | B1205 |
| CellEvent$^{TM}$ Caspase-3/7 Green Detection Reagent | ThermoFisher Scientific | C10423 |
| Live/Dead BacLight viability kit for microscopy | ThermoFisher Scientific | L7007 |
| XTT (sodium 3´-[1- (phenylaminocarbonyl)- 3,4-tetrazolium]-bis (4-methoxy6-nitro) benzene sulfonic acid hydrate | ThermoFisher Scientific | X6493 |
| lipofectamine 3000 | ThermoFisher Scientific | L3000075 |
| Fluo-8-AM | Abcam | ab142773 |
| Normal Donkey Serum | Abcam | ab7475 |
| *Commercial Assays* | | |
| TG1 ELISA | Aviva Systems Bio | OKCD01601 |
| *Cell Lines* | | |
| 16HBE | Dieter Gruenert Lab, UCSF [9] | Available through Dr. Beate Illek, Children's Hospital Oakland |
| Beas-2B | ATCC | CRL-9609 |
| NCI-H292 | ATCC | CRL-1848 |
| Caco-2 | ATCC | HTB-37 |
| A549 | ATCC | CCL-185 |
| HEK293T | ATCC | CRL-11268 |
| Calu-3 | ATCC | HTB-55 |
| RPMI 2650 | ATCC | CCL-30 |
| T2R39 knockout A549s | This study | N/A |
| Primary human bronchial epithelial cells (HBEs) | Lonza | CC-2450 |
| Primary nasal epithelial cells | This study | N/A |
| ELE2 hBMI immortalized bronchial epithelial cells | This study | N/A |
| Z0E hBMI immortalized nasal epithelial cells | This study | N/A |
| *Primers for qPCR* | | |
| TaqMan Primers for T2R1 | ThermoFisher Scientific | Hs00251930_s1 |
| TaqMan Primers for T2R4 | ThermoFisher Scientific | Hs00249946_s1 |
| TaqMan Primers for T2R7 | ThermoFisher Scientific | Hs00256778_s1 |

| | | |
|---|---|---|
| TaqMan Primers for T2R8 | ThermoFisher Scientific | Hs00256766_s1 |
| TaqMan Primers for T2R9 | ThermoFisher Scientific | Hs00256757_s1 |
| TaqMan Primers for T2R10 | ThermoFisher Scientific | Hs00256794_s1 |
| TaqMan Primers for T2R13 | ThermoFisher Scientific | Hs00256781_s1 |
| TaqMan Primers for T2R14 | ThermoFisher Scientific | Hs00256800_s1 |
| TaqMan Primers for T2R16 | ThermoFisher Scientific | Hs00249955_s1 |
| TaqMan Primers for T2R20 | ThermoFisher Scientific | Hs00604340_s1 |
| TaqMan Primers for T2R30 | ThermoFisher Scientific | Hs03054740_sH |
| TaqMan Primers for T2R31 | ThermoFisher Scientific | Hs00604313_sH |
| TaqMan Primers for T2R38 | ThermoFisher Scientific | Hs00604294_s1 |
| TaqMan Primers for T2R39 | ThermoFisher Scientific | Hs00603443_s1 |
| TaqMan Primers for T2R40 | ThermoFisher Scientific | Hs00602589_s1 |
| TaqMan Primers for T2R43 | ThermoFisher Scientific | Hs00853105_sH |
| TaqMan Primers for T2R46 | ThermoFisher Scientific | Hs00853124_s1 |
| TaqMan Primers for UBC | ThermoFisher Scientific | Hs01871556_s1 |
| *Recombinant DNA* | | |
| nls-R-GECO | Addgene | 32462 |
| G-GECO1.2 | Addgene | 32446 |
| D1ER | Addgene | 36325 |
| 4mtD3cpv | Addgene | 36324 |
| Flamindo2 | Addgene | 73938 |
| nls-Flamindo2 | Addgene | 73939 |
| GFP T2R14 promoter construct; pEZX ProTAS2R14, created for this study; Promoter of TAS2R14 NM_023922 | Genecopoeia | N/A |
| hBMI lentiviral vector (pLV[Exp]-CMV>hBMI1[NM_005180.8]) | VB161220-1120wck | VB170413-1041avf |
| pRS TAS2R8 shRNA plasmid | OriGene | T301219B |
| pRS T2R10 shRNA plasmid | OriGene | T301240B |
| pRS T2R14 shRNA plasmid | OriGene | TR301238A |
| pRS T2R14 shRNA plasmid | OriGene | TR301238B |
| pRS scramble shRNA plasmid | OriGene | TR30012 |
| myc-T2R14 (pRP[Exp] CMV>Myc/hTAS2R14[NM_023922.1]*) | Cyagen Biosciences | VB170413-1041avf |
| myc-T2R39 (pRP[Exp] CMV>Myc/hTAS2R39[NM_176881.2]*) | Cyagen Biosciences | VB170424-1092bmx |
| Myc-T2R10 (pRP[Exp] CMV>Myc/hTAS2R10[NM_023921.1]*) | Cyagen Biosciences | VB170412-1090rje |
| GFP-T2R39 (pRP[Exp] CMV>EGFP(ns):3XGGGGS:hTAS2R39[NM_176881.2]*) | Cyagen Biosciences | VB170424-1098rxn |
| T2R39-GFP (pRP[Exp] CMV>hTAS2R39[NM_176881.2](ns):3XGGGGS:EGFP*) | Cyagen Biosciences | VB170424-1112ayf |
| Control Lentiviral CRISPR knockout vector | Cyagen Biosciences | VB161020-1072gqy |

| | | |
|---|---|---|
| T2R39 Lentiviral CRISPR knockout vector (pLV[CRISPR]-hCAS9:T2A:Hygro-U6>hTAS2R39_1_458_20nt:U6>hTAS2R39_2_606_20nt | Cyagen Biosciences | VB161020-1082cvg |
| *Software* | | |
| MetaFluor | Molecular Devices | N/A |
| MetaMorph | Molecular Devices | N/A |
| QuantStudio 5 | Applied Biosystems, Inc | N/A |
| Prism v8 | GraphPad Software | |
| ImageJ/FIJI | Open Source [13] | N/A |

### Supplementary Table S2: T2R agonists used in this study and associated receptors previously identified by heterologous expression studies.

| **Compound** | **Human T2Rs activated (EC[1] in μM)** | **References** |
|---|---|---|
| Apigenin | T2R14 (8), T2R39 (1) | [1, 14, 15, 16] |
| Denatonium benzoate | T2R4 (300), T2R8 (1000), T2R10 (3), T2R13 (30), T2R39 (100), T2R43 (300), T2R46 (30), T2R47 (0.03) | [15, 16] |
| Diphenidol | T2R1 (100), T2R4 (100), TR7 (10), T2R10 (30), T2R13 (30), T2R14 (10), T2R16 (100), T2R38 (100), T2R39 (100), T2R40 (30), T2R43 (30), T2R44 (3), T2R46 (30), T2R47 (100), T2R49 (100) | [15, 16] |
| Diphenhydramine (DPD) | T2R14 (30), T2R40 (30) | [15, 16] |
| Flufenamic acid (FFA) | T2R14 (0.1) | [15, 16] |
| 3-oxo-C12HSL[2,3] | T2R4 ($EC_{50}$ = 41), T2R10 (ND), T2R14 ($EC_{50}$ = 70), T2R20 ($EC_{50}$ = 58), T2R38 (ND) | [4, 15, 17, 18] |
| Parthenolide | T2R1 (100), T2R4 (30), T2R8 (100), T2R10 (30), T2R14 (3), T2R44 (100), T2R46 (1). | [15, 16] |
| Phenylthiocarbamine (PTC) | T2R38 (0.04) | [15, 16] |
| Quinine | T2R4 (10), T2R7 (10), T2R10 (10), T2R14 (10), T2R39 (10), T2R40 (10), T2R43 (10), T2R44 (10), T2R46 (10) | [15, 16] |
| Sodium Benzoate | T2R14 (3000), T2R16 (300) | [15, 16] |
| (-)α-Thujone | T2R10 (100), T2R14 (3) | [15, 16] |

[1]Effective concentration (EC) is defined as the minimal concentration of agonist that elicits a detectable response, largely based on *in vitro* heterologous expression assays. $EC_{50}$, defined as the concentration of agonist that elicits a response 50% of the maximal response, is used when indicated
[2]"ND" denotes EC not determined
[3]Not tested against all 25 human T2Rs

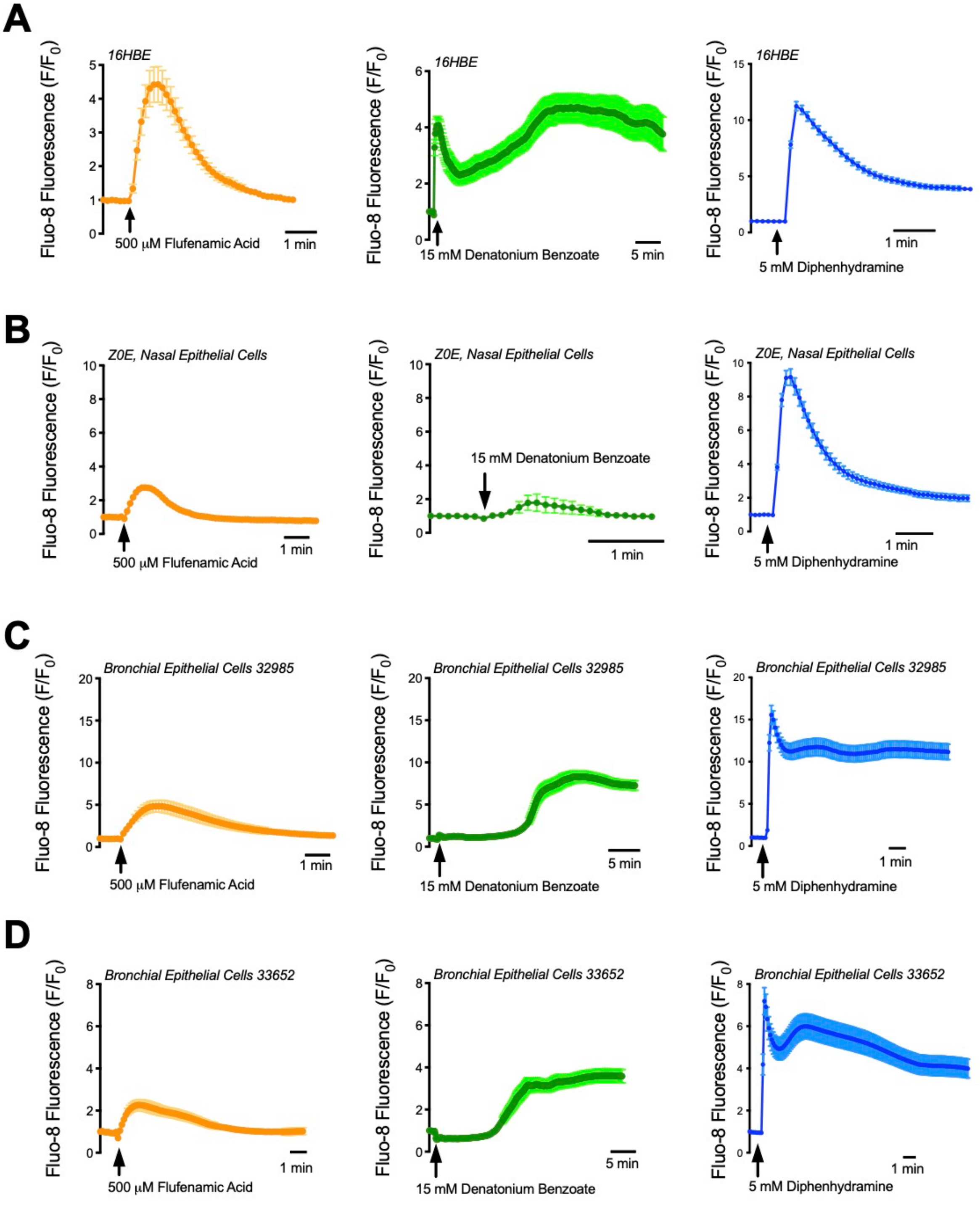


**Supplementary Figure S1. T2R agonists activate $Ca^{2+}_i$ release in cultured airway epithelial cells. (A)** 16HBE cells and **(B-D)** undifferentiated primary airway epithelial cells pre-incubated with $Ca^{2+}$ binding dye, Fluo-8 AM, release $Ca^{2+}_i$ in response to T2R agonists diphenhydramine, flufenamic acid, and denatonium benzoate. For all images one representative trace from 3-4 experiments are shown. Data points on bar graphs represent the mean ± SEM from 3-4 experiments.

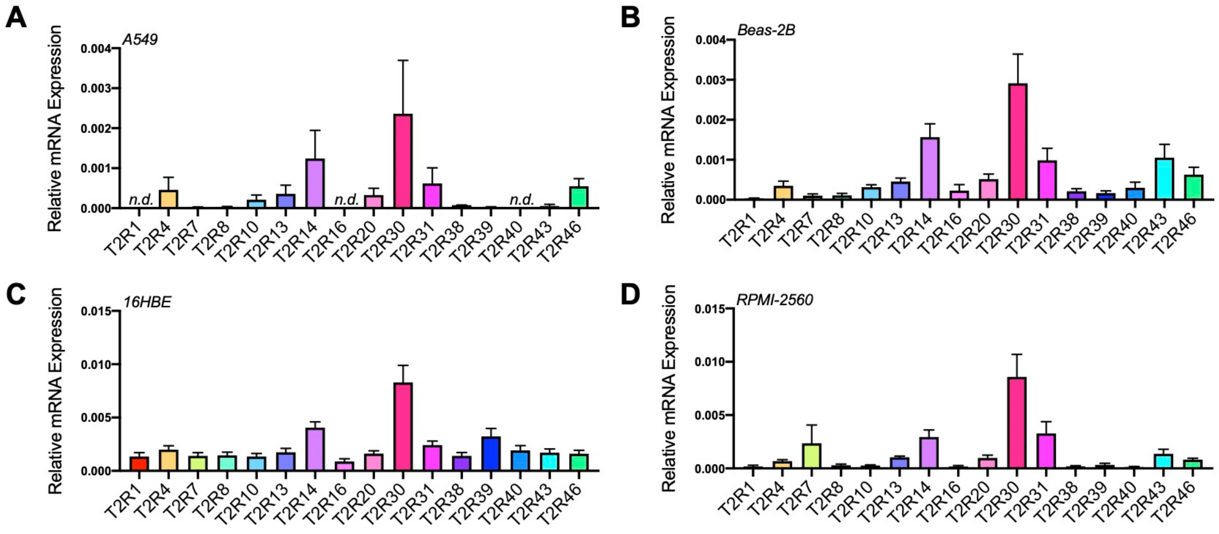


**Supplementary Fig. S2 Characterization of T2R mRNA Expression in Airway Cell Lines.** **A-D** Messenger RNA expression of various T2Rs (relative to Ubiquitin C [UBC]) of airway cell lines A549 (*A*), Beas-2B (*B*), 16HBE (*C*), and RPMI-2650 (*D*). Briefly, cultures were suspended in TRIzol then RNA was isolated via spin column purification. After quantification, 400 ng of RNA was used as a template for RT-PCR, after which the resulting cDNA was diluted so that each qPCR reaction contained approximately 4 ng of cDNA.

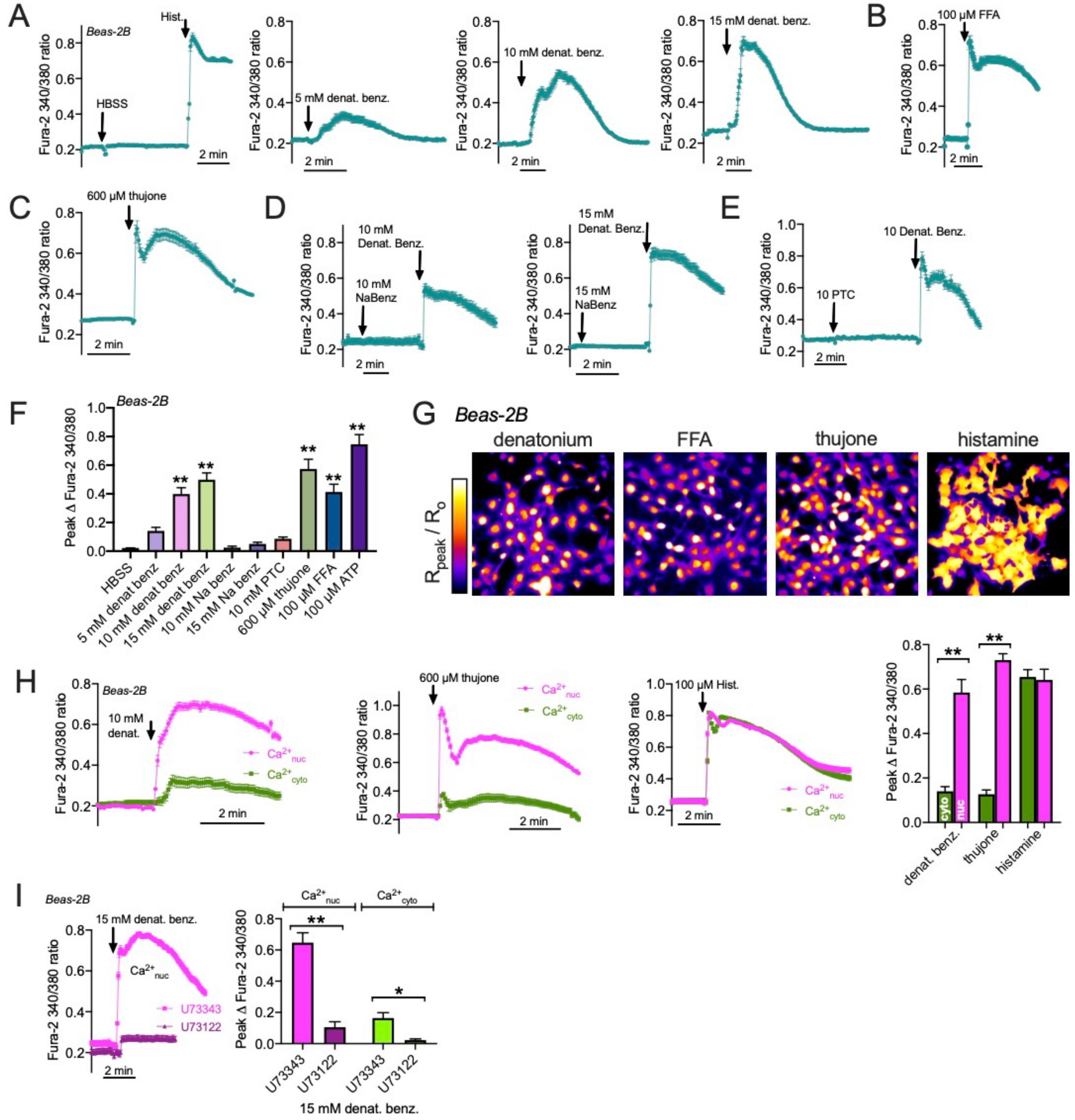


**Supplementary Fig. S3 $Ca^{2+}$ responses in Beas-2B cells loaded with fura-2 appear to be largely nuclear in response to T2R agonists but not histamine. A** Representative $Ca^{2+}$ traces (approximated by fura-2 340/380 excitation ratio; R) from individual experiments (mean ± SEM: n = 22 to 41 cells per experiment) of Beas-2B cells loaded with fura-2 on 8 well chamber slides (45 min; 5 μM in HBSS). Cells were stimulated with HBSS (control; pipetted into the well the same as HBSS with dissolved agonists), histamine (100 μM), or denatonium benzoate (5, 10, or 15 mM; multi-T2R agonist) dissolved directly into HBSS. Shown are global $Ca^{2+}$ traces from circling the entire cell. **B-C** $Ca^{2+}$ trace as in *a*, but cells were stimulated with 100 μM FFA (*b*; T2R14 agonist) or 600 µM thujone (*c*; T2R10 and T2R14 agonist). **D-E** Representative experiments showing lack of response to sodium benzoate (NaBenzoate; 15 mM; *d*) or PTC (*e*; T2R38 agonist; 10 mM) in cells with intact response to equal concentration of denatonium benzoate. **F** Bar graph of peak responses from independent experiments (n = 4-10) as shown in (A-E). Significance determined by 1-way ANOVA with Dunnett's posttest comparing all values to control (addition of HBSS only); $^{**}p<0.01$. **G** Representative images of fura-2 peak 340/380 ratio ($R_{peak}$) divided by initial 340/380 ratio ($R_o$) showing nuclear appearance of responses to denatonium, FFA, and thujone but more uniform cellular response to histamine. 340/380 ratio images ($R_o$ and $R_{peak}$) were first created as 32-bit floating-point images from background-subtracted raw 340 and 380 images using the image calculator and then similarly divided to create $R_{peak}/R_o$ image. **H** Nuclear regions of interest were estimated by thresholding $R_{peak}/R_o$ images using ImageJ/FIJI and used measure $Ca^{2+}$ changes within the apparent nuclear region ($Ca^{2+}_{nuc;}$ shown in magenta). The nuclear ROIs were also used to create a binary mask, and apparent cytosolic $Ca^{2+}$ ($Ca^{2+}_{cyto;}$ shown in green) was approximated by taking the cellular 340/380 ratio without the identified nuclear region. The traces shown are mean ± SEM of representative experiments (~30 cells per experiment) as shown in (A-C) but with $Ca^{2+}_{nuc}$ vs $Ca^{2+}_{cyto}$ separated out. Bar graph is mean ± SEM of 6 independent experiments analyzed in this way. Significance determined by one-way ANOVA with Bonferroni posttest comparing $Ca^{2+}_{nuc}$ vs $Ca^{2+}_{cyto}$ for each agonist; $^{**}p<0.01$. **I** Nuclear vs cytosolic $Ca^{2+}$ responses analyzed as in (H) were measured in cells stimulated after 45 min pretreatment with 10 µM U73122 (PLC inhibitor) or inactive analogue U73122. Significance determined by one-way ANOVA with Bonferroni posttest comparing U73343 vs U73122 each for $Ca^{2+}_{nuc}$ and $Ca^{2+}_{cyto}$ separately; $^{**}p<0.01$

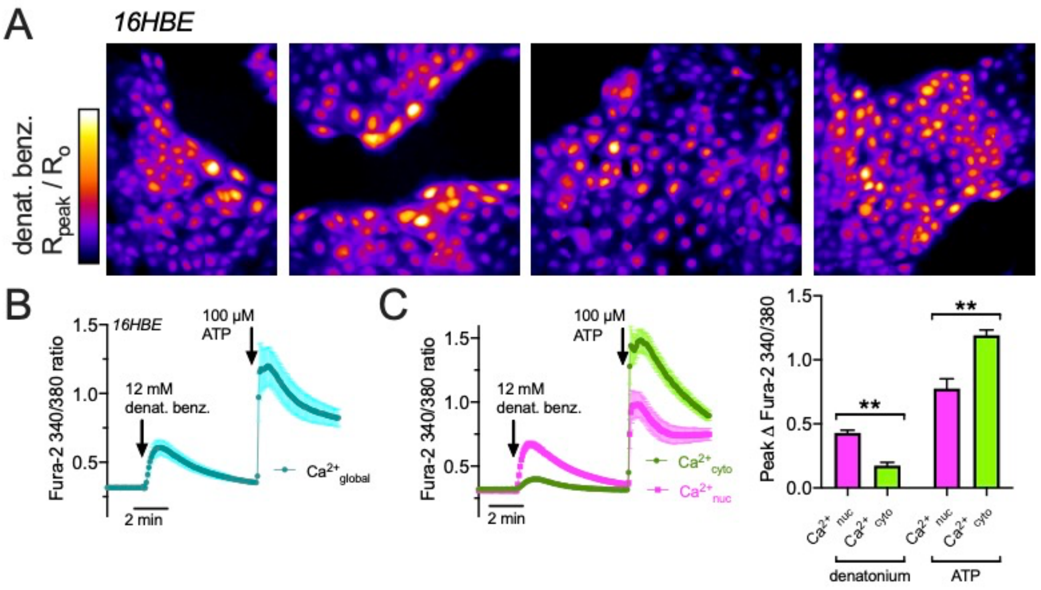


**Supplementary Fig. S4  $Ca^{2+}$ responses in 16HBE cells loaded with Fura-2 appear to be largely nuclear in response to T2R agonists but not ATP.  A** Representative $R_{peak}/R_o$ images in 16HBE cells loaded with fura-2 and stimulated with 12 mM denatonium benzoate as shown for Beas-2B cells in Figure S2G showing nuclear appearance of $Ca^{2+}$ responses. **B-C** Representative trace from one experiment with cells (n = 12) stimulated with denatonium followed by 100 μM ATP, analyzed as in Figure S2H.  Global $Ca^{2+}$ responses ($Ca^{2+}$ global) are shown in black in (B), while separated $Ca^{2+}_{nuc}$ vs $Ca^{2+}_{cyto}$ are shown in (C).  Bar graph in (C) shows mean ± SEM of peak $Ca^{2+}_{nuc}$ vs $Ca^{2+}_{cyto}$ for denatonium benzoate and ATP stimulation from 5 independent experiments.  Significance determined by one-way ANOVA with Bonferroni posttest comparing $Ca^{2+}_{nuc}$ vs $Ca^{2+}_{cyto}$ for each agonist; ***p*<0.01.

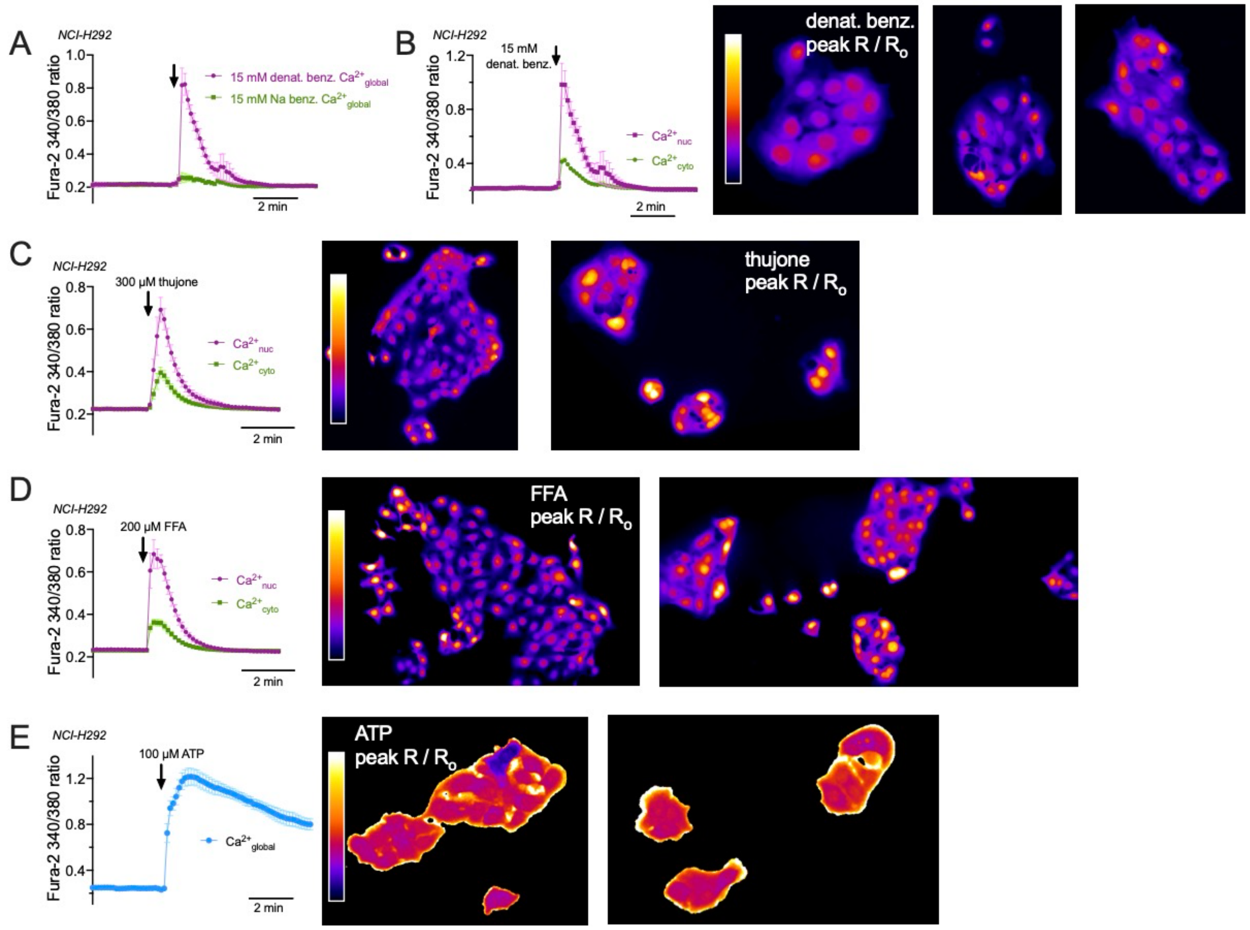


**Supplementary Fig. S5 $Ca^{2+}$ responses in NCI-H292 cells loaded with Fura-2 appear to be largely nuclear in response to T2R agonists but not ATP. A** Average trace from 6 independent experiments of H292 cells stimulated with 15 mM denatonium benzoate (magenta) or sodium benzoate (NaBenzoate; green) showing $Ca^{2+}$ response to denatonium benzoate but not equivalent concentration of sodium benzoate. **B-D** Representative traces (analyzed as in Figure S2H) and peak $R/R_o$ images (as constructed for Beas-2B cells in Figure S2G), in NCI-H292 cells loaded with fura-2 and stimulated with denatonium benzoate (B), thujone (C), or FFA (D) showing apparent nuclear $Ca^{2+}$ responses to T2R agonist stimulation. **E** trace of global $Ca^{2+}$ response to ATP and peak R/Ro image showing more uniform elevation of $Ca^{2+}$.

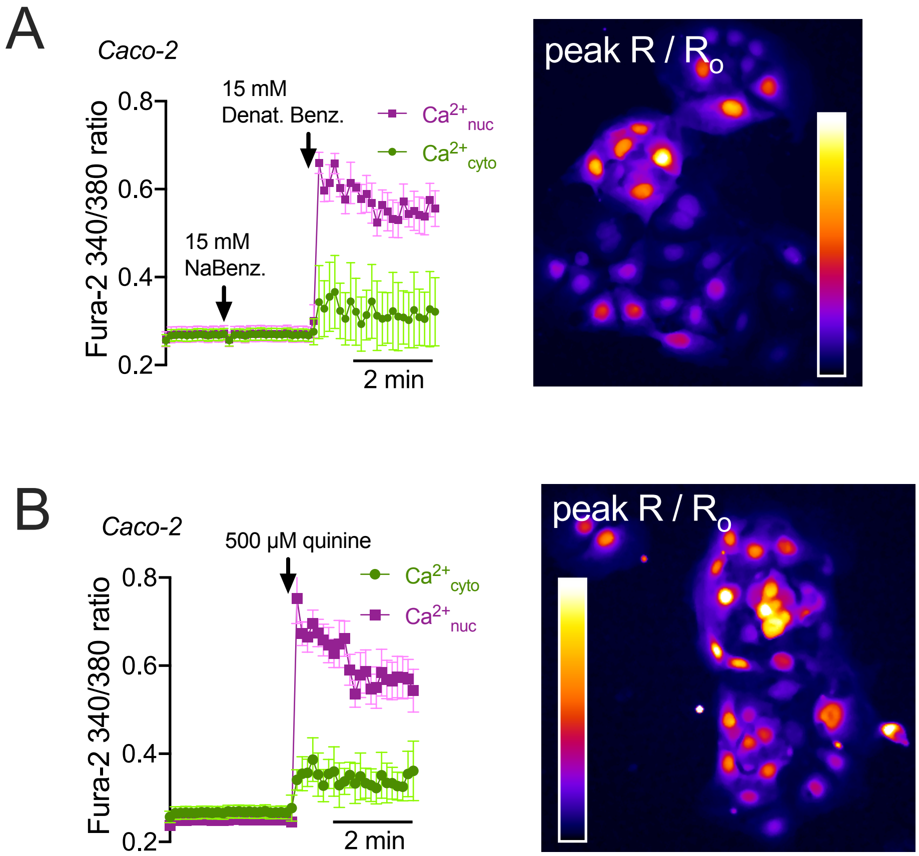


**Supplementary Fig. S6 $Ca^{2+}$ responses in Caco-2 cells loaded with Fura-2 appear to be largely nuclear in response to T2R agonists. A-B** Representative traces (analyzed as in Figure S2H) and peak R/$R_o$ images (as constructed for Beas-2B cells in Figure S2G) showing apparent nuclear $Ca^{2+}$ responses to denatonium benzoate (A) or quinine sulfate (B). Traces are representative data from 3 experiments.

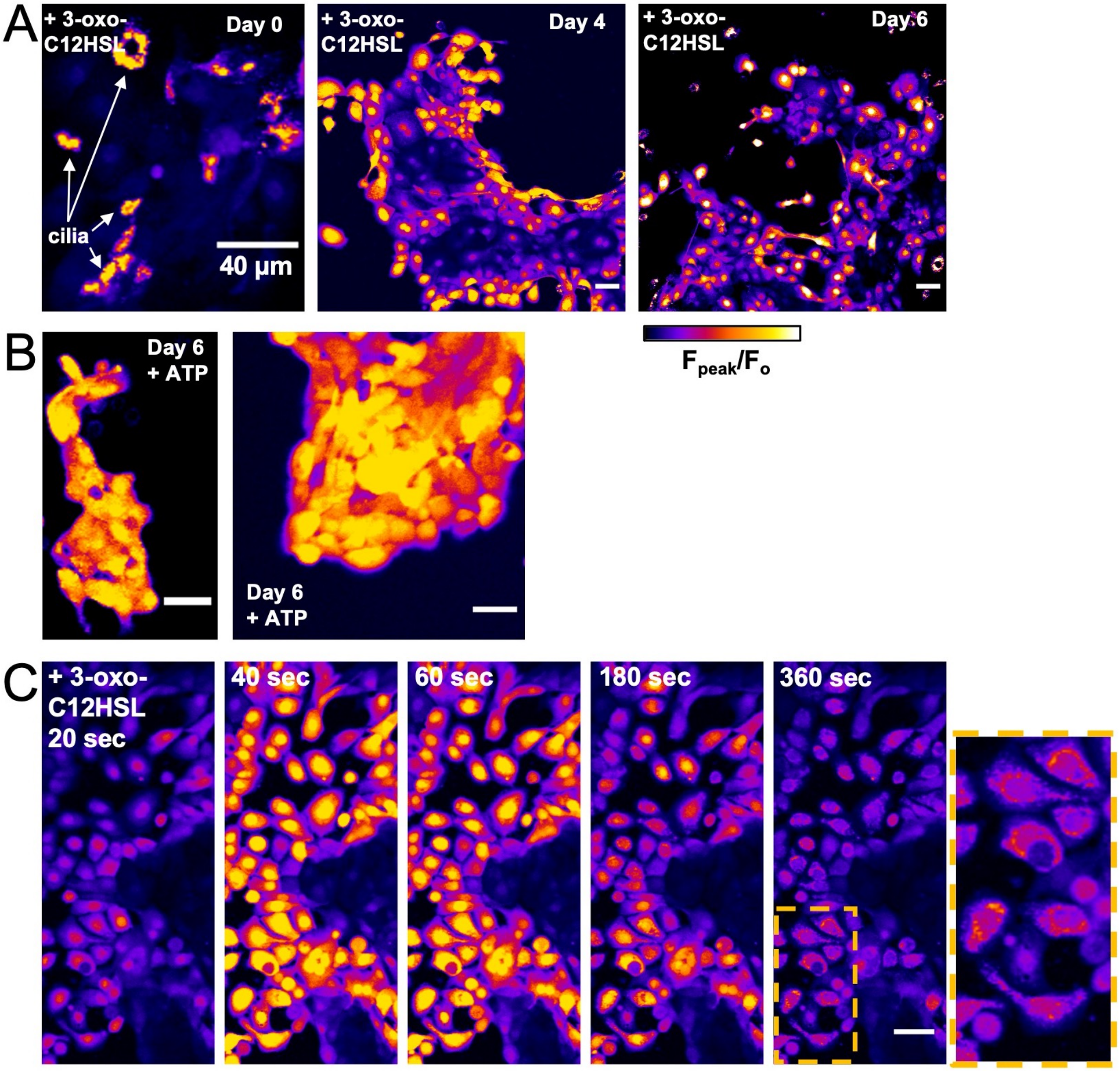


**Supplementary Fig. S7 $Ca^{2+}$ responses to 3-oxo-C12HSL in primary nasal cells loaded with Fluo-4 appear to shift from cilia to nuclear after 4-6 days in culture. A** $F_{peak}/F_o$ images of peak responses to 100 µM 3-oxo-C12HSL in ciliated cells removed from nasal brushings (day 0) and then cultured in submersion (Lonza BEBM basal media plus SingleQuot growth supplements) on plastic over 4-6 days (during which time, cilia are lost and spread out into a more squamous morphology). Note the change in response from cilia-localized at day 0 to nuclear localized at days 4-6. The intensity of images first three images are scaled identically, thus the magnitude of the cilia-localized response was similar to the nuclear-localized response. **B** Responses to ATP at day 6, which elicited a more global cellular $Ca^{2+}$ responses. **C** Fluo-4 responses to 3-oxo-C12HSL in cultured submerged cells (day 6 shown) appeared to initiate in the nucleus and resulted in more sustained $Ca^{2+}$ elevation in the perinuclear region, possibly the mitochondria. All images are representative of results observed from cells isolated and cultured from 3 different patent *TAS2R38* PAV/PAV patient turbinates. All scale bars are 40 µm.

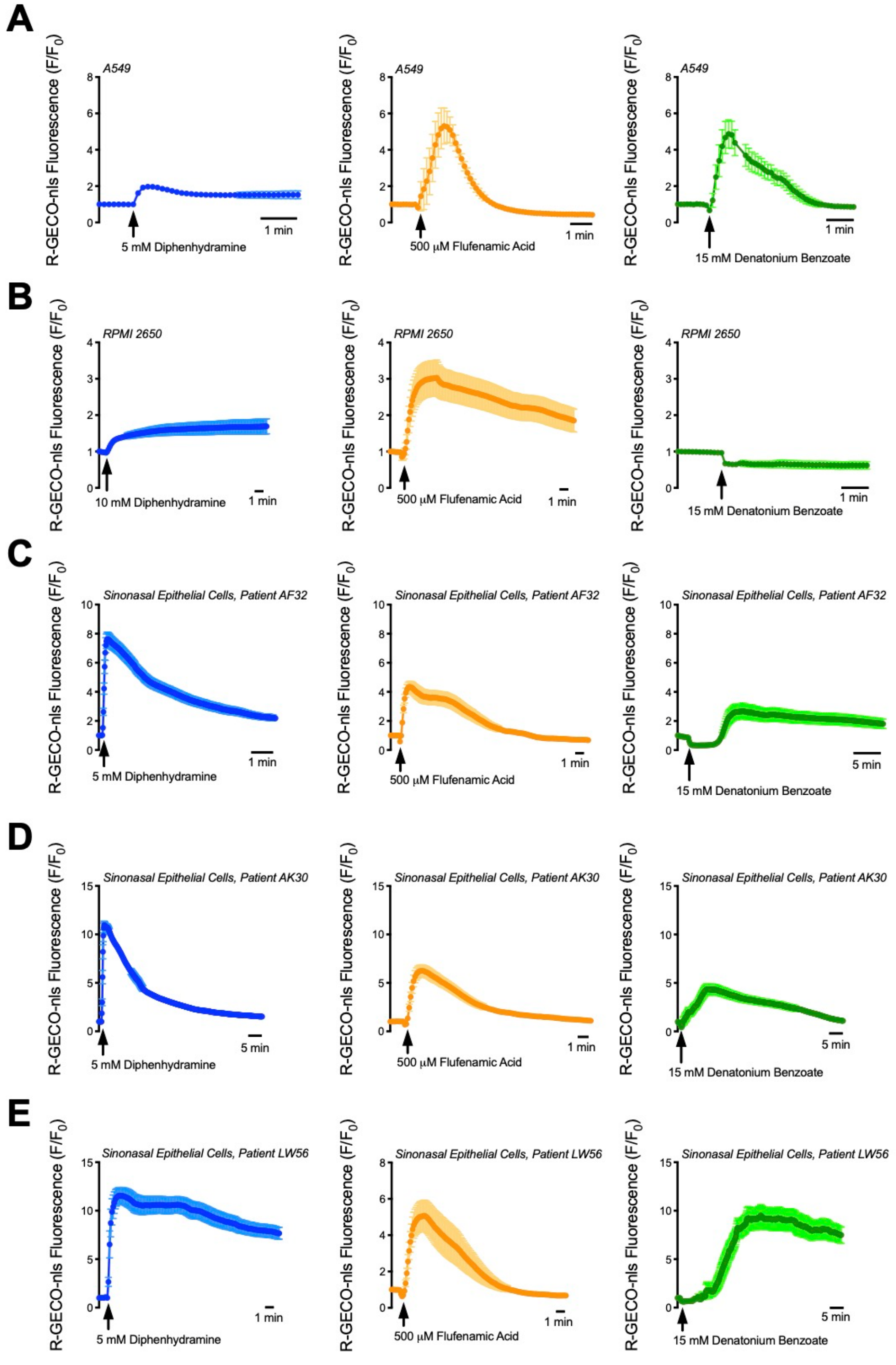


**Supplementary Fig. S8 T2R agonists release nuclear calcium in airway cell lines and primary sinonasal epithelial cells. A-E** Airway cell lines (A-B) and primary nasal cells from three patients (C-E) were transfected with R-GECO-nls 48 hours prior to the experiment [10]. All airway cells tested, with exception of RPMI 2650 treated with denatonium benzoate, release $Ca^{2+}_{nuc}$ in response to a variety of agonists that interact with a wide range of T2R's (as seen in Table S1). For all data 1 representative trace from ≥3 experiments shown.

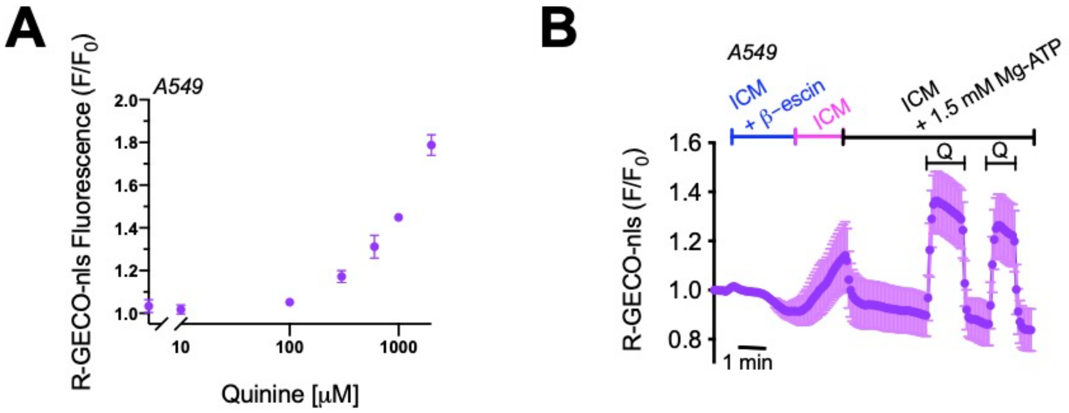


**Supplementary Fig. S9 Activation of $Ca^{2+}_{nuc}$ responses in permeabilized A549 cells. A** Quinine elevates $Ca^{2+}_{nuc}$ in intact A549 cells in a dose-dependent manner. **B** A549 cells expressing R-GECO-nls were briefly permeabilized with β-escin in intracellular-like media (ICM) plus ATP to operate $Ca^{2+}$ pumps and perfused 670 μM quinine. Note that ATP activates a decrease in $Ca^{2+}$ (suggesting pump-operated $Ca^{2+}$ uptake into stores) rather than an increase in $Ca^{2+}$ (suggesting plasma membrane permeabilization has inactivated plasma membrane purinergic receptors). This suggests T2Rs are functional on nuclear membranes and can increase $Ca^{2+}_{nuc}$. All data are representative of ≥3 independent experiments.

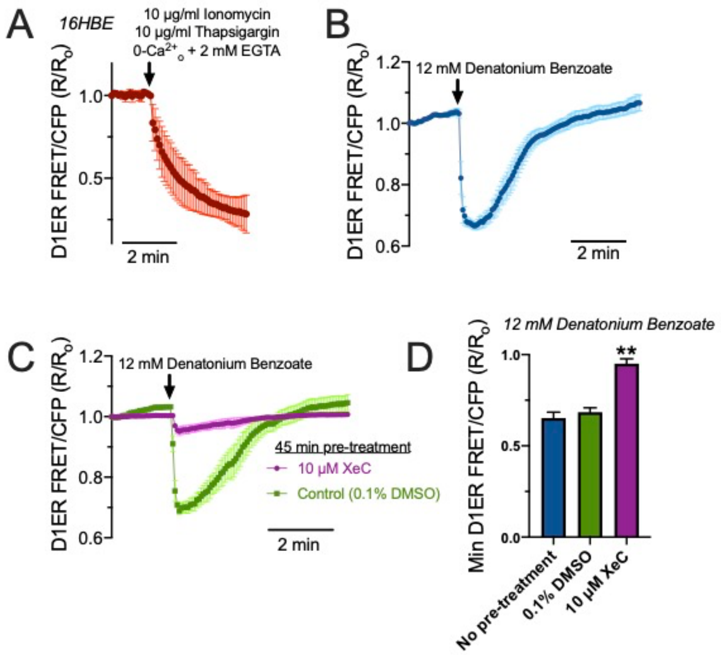


**Supplementary Fig. S10 Release of $Ca^{2+}$ from ER stores during denatonium benzoate stimulation in 16HBE cells.** $Ca^{2+}_{nuc}$ signaling is likely distinct from cytosolic $Ca^{2+}$ signaling, though likely regulated by similar pathways; in the nucleus, $Ca^{2+}$ elevations can be initiated through either nuclear $IP_3$-dependent pathways or ryanodine receptors [19, 20, 21, 22, 23, 24, 25, 26, 27]. In some early studies, increases in $Ca^{2+}_{nuc}$ were thought to be tightly and passively tied to cytosolic $Ca^{2+}$ [28], while more recent studies suggest they can be independently regulated [19, 29, 30]. $Ca^{2+}$ microdomains may be created by restricted diffusion across the nuclear membrane or nuclear pore complexes and/or $Ca^{2+}$ buffering from the nucleoplasmic reticulum or other organelles like ER or mitochondria [25, 31]. The nuclear envelope is continuous with the endoplasmic reticulum, containing both $IP_3$Rs and RyRs on the inner nuclear membrane. Thus, the nuclear envelope is itself is a $Ca^{2+}$ store capable of releasing $Ca^{2+}$ into the nucleus [32, 33, 34]. Invaginations, tubules, or sheets of inner nuclear envelope membrane can form a nucleoplasmic reticulum in many cells that also serves as an intranuclear $Ca^{2+}$ store and signaling hub [20, 35, 36]. Small (~50 nm) membrane-bound vesicles in the nucleoplasm may also contain $IP_3$ receptors and facilitate nuclear GPCR $Ca^{2+}$ signaling [37]. We wanted to test if T2R stimulation activated bulk $Ca^{2+}$ depletion from the ER or if release was occuring from more local sources (e.g., nuceloplasmic reticulum granules). To do this, 16HBE cells were transfected with ratiometric CFP/YFP FRET-based ER $Ca^{2+}$ indicator D1ER [38] and imaged 24 hours later. **A** Stimulation of cells with $Ca^{2+}$ ionophore ionomycin and $Ca^{2+}$ ATPase inhibitor thaspigargin in the absence of extracellular $Ca^{2+}$ (0-$Ca^{2+}_o$ HBSS; no added $Ca^{2+}$ plus 2 mM EGTA) to fully deplete ER $Ca^{2+}$ stores and show the dynamic range of the indictor. Shown is average of 5 transfected cells from 5 independent experiments imaged at 60x. **B** Stimulation with 12 mM denatonium benzoate resulted in transient reduction of ER $Ca^{2+}$. Shown is average of 4 transfected cells from 4 independent experiments imaged at 60x. **C** Denatonium benzoate-depletion of ER $Ca^{2+}$ was blocked by 45 min pre-incubation by $IP_3$R inhibitor xestospongin C (XeC) but not by vehicle control (0.1% DMSO). Shown is average of 4 transfected cells from 4 independent experiments imaged at 60x. **D** Min D1ER FRET/CFP $R/R_o$ values were plotted from experiments shown in (B) and (C). Significance determined by one-way ANOVA with Bonferroni posttest; ***p*<0.01 vs either no pretreatment or 0.1% DMSO vehicle control. Our data suggest that the elevations in $Ca^{2+}_{nuc}$, at least in part, originate from the bulk cellular ER $Ca^{2+}$ stores.

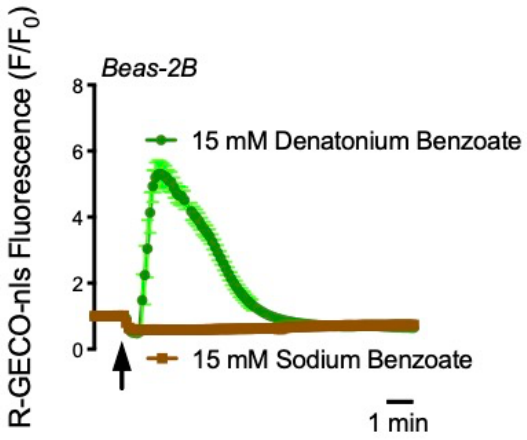


**Supplementary Fig. S11 Denatonium benzoate but not sodium benzoate activated $Ca^{2+}_{nuc}$ in Beas-2B cells.** Shown are two traces of R-GECO-nls fluorescence. Traces are representative of ≥3 independent experiments.

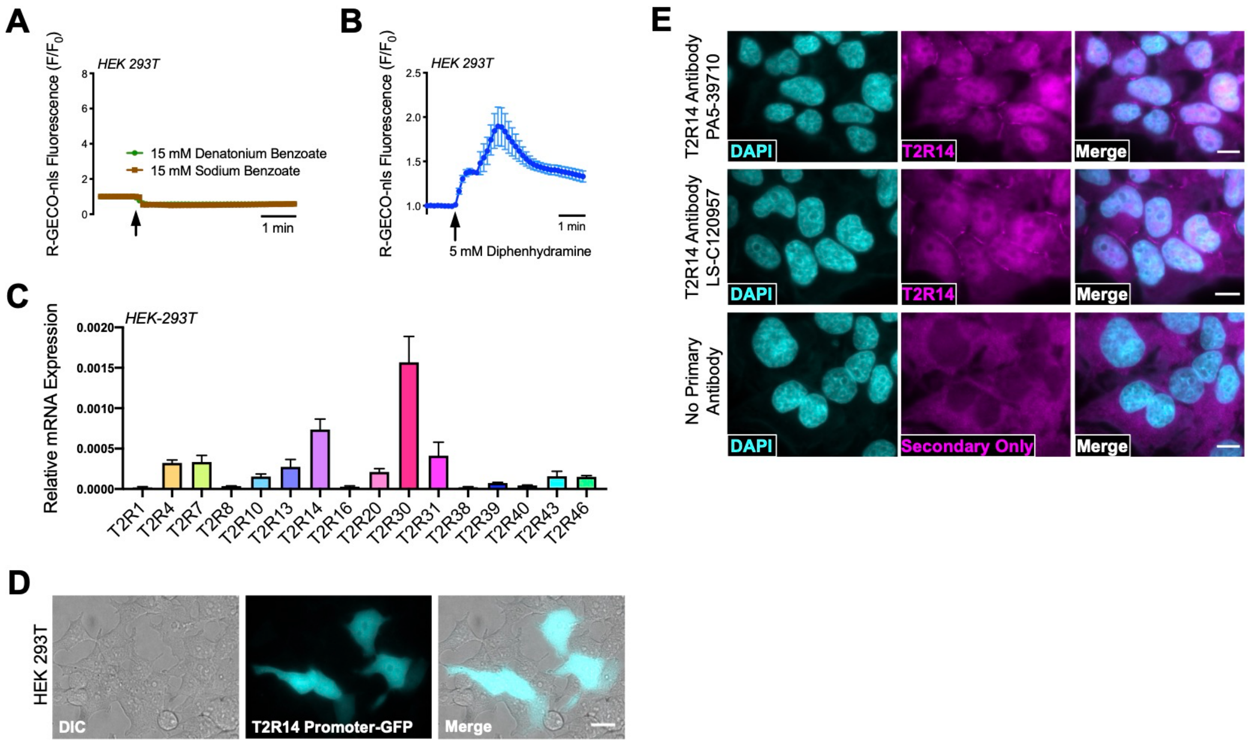


**Supplementary Fig. S12 HEK293T cells express endogenous T2R14. A-B** HEK cells were unresponsive to denatonium (not a T2R14 agonist) or sodium benzoate (very low affinity T2R14 agonist). However, T2R14 agonist diphenhydramine (~100 fold greater affinity for T2R14 than sodium benzoate) caused a $Ca^{2+}_{nuc}$ release. **C** Messenger RNA expression of various T2Rs (relative to Ubiquitin C) of HEK-293T **D** HEK's were transfected with a construct containing 1,452 bp upstream region of the *TAS2R14* gene leading into the EGFP gene 72 hours prior to visualization via fluorescent microscopy. Scale bar represents 20 µm. **E** Immunofluorescence showing plasma membrane localization and possibly some nuclear localization of T2R14 in HEK293Ts. These data confirm that HEKs express endogenous functional T2Rs. The lower threshold for activation of $Ca^{2+}$ responses by bitter compounds observed after transfection of tagged T2Rs may be due partly to an increase in receptor numbers or may also be due to an increase in plasma membrane localization due to construct tagging. We suggest caution when using HEK cells as a T2R expression model.

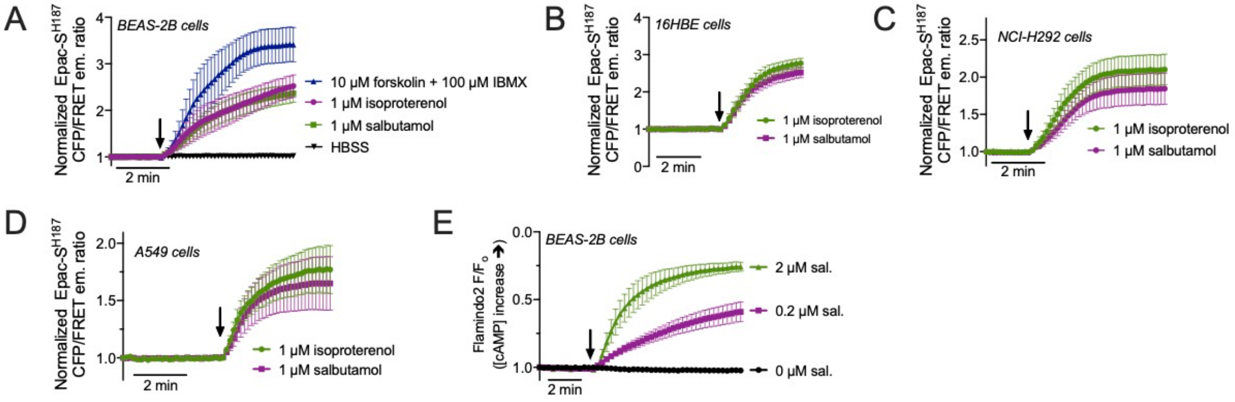


**Supplementary Fig. S13 cAMP responses to β2AR agonists. A-C** Beas-2Bs (*A*), 16HBEs (*B*), H292s (*C*), or A549s (*D*) were transfected with EPAC-SH$^{187}$ [39] and Flamindo2 with and imaged during stimulation with β2AR agonists isoproterenol or salbumatol. Upward deflection indicates increase in cAMP. **D** Beas-2Bs were transfected with Flamindo and imaged during stimulation with isoproterenol or salbumatol. Experiments are representative traces of ≥3 independent experiments.

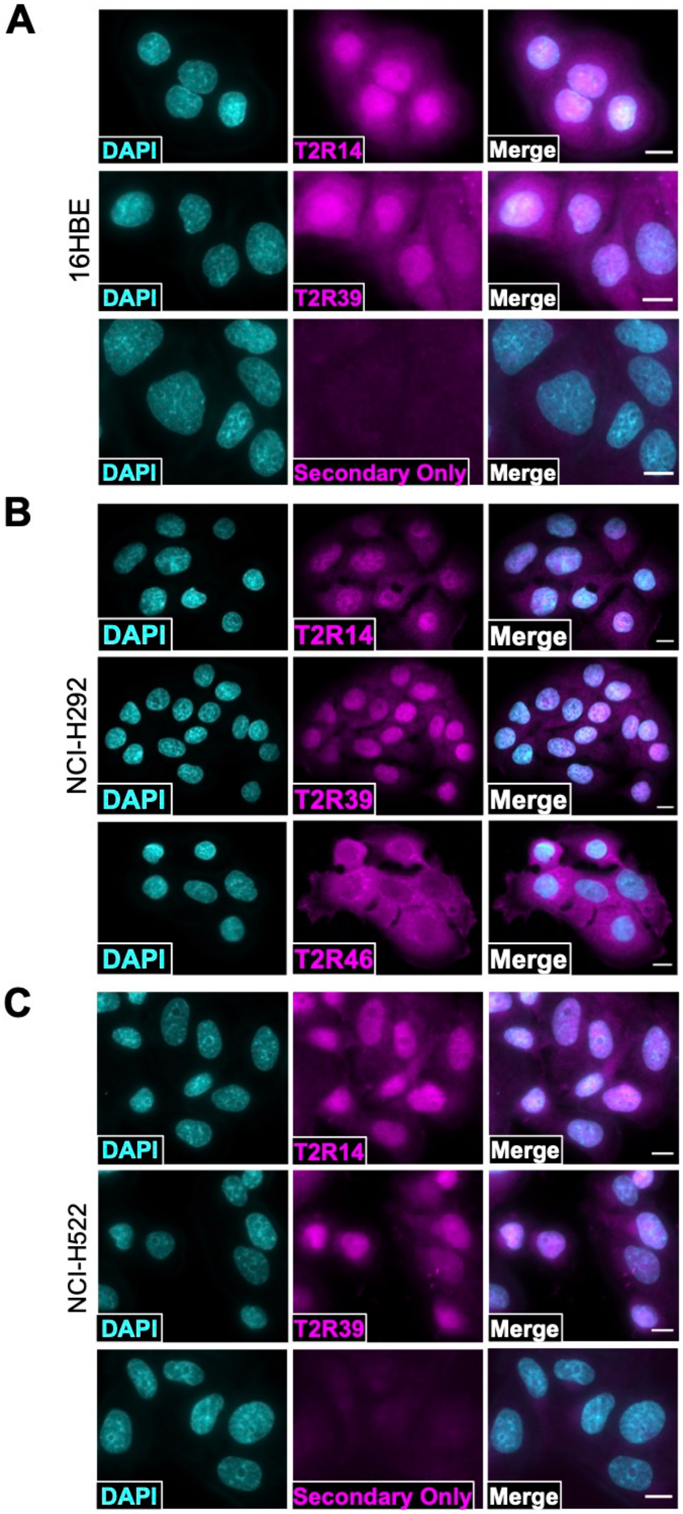


**Supplementary Fig. S14 Nuclear localization of T2R14 and 39 A-C** Immunofluorescence images showing T2R14 and T2R39 localization in 16HBE (*A*), H292 (*B*), and H522 (*C*) cells. Note T2R46 was not nuclear localized in H292s (*B*). For all images, scale bars represent 10 µm.

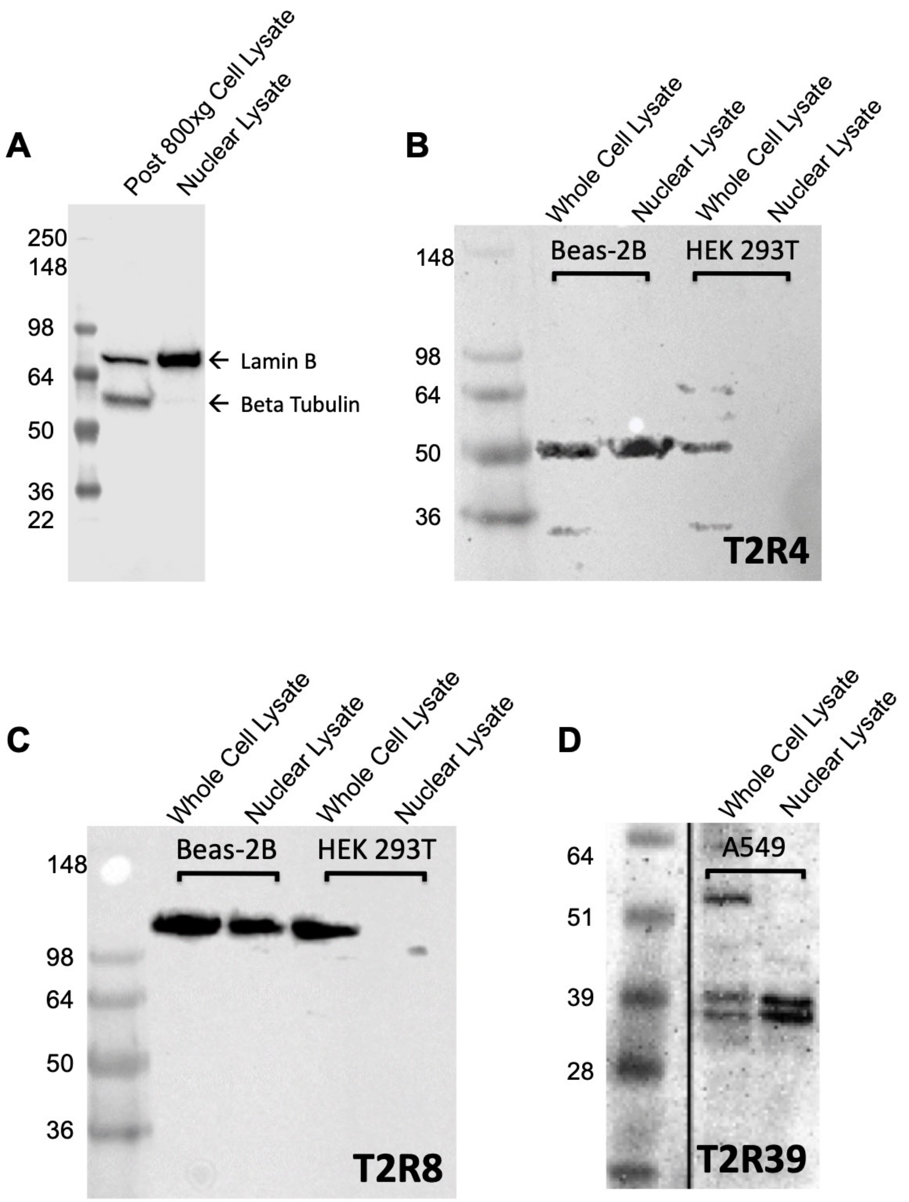


**Supplementary Fig. S15 T2R4, T2R8, and T2R39 are detectable in nuclear-enriched cell lysates in airway cells via Western Blot.** **A** Cell lysates are enriched for nuclear extract utilizing the REAP method [12] as denoted by nuclear marker Lamin B and cytosolic marker Beta Tubulin. **B-C** Images of immunoblots revealing T2R4 and T2R8 expression in post 800 x *g* lysates vs. nuclear enriched lysates from Beas-2B and HEK cultures. **D** T2R39 enrichment in A549 nuclear extracts. All images are representative of 3 experiments; in all protein gels, 60 µg of protein extract was loaded per lane.

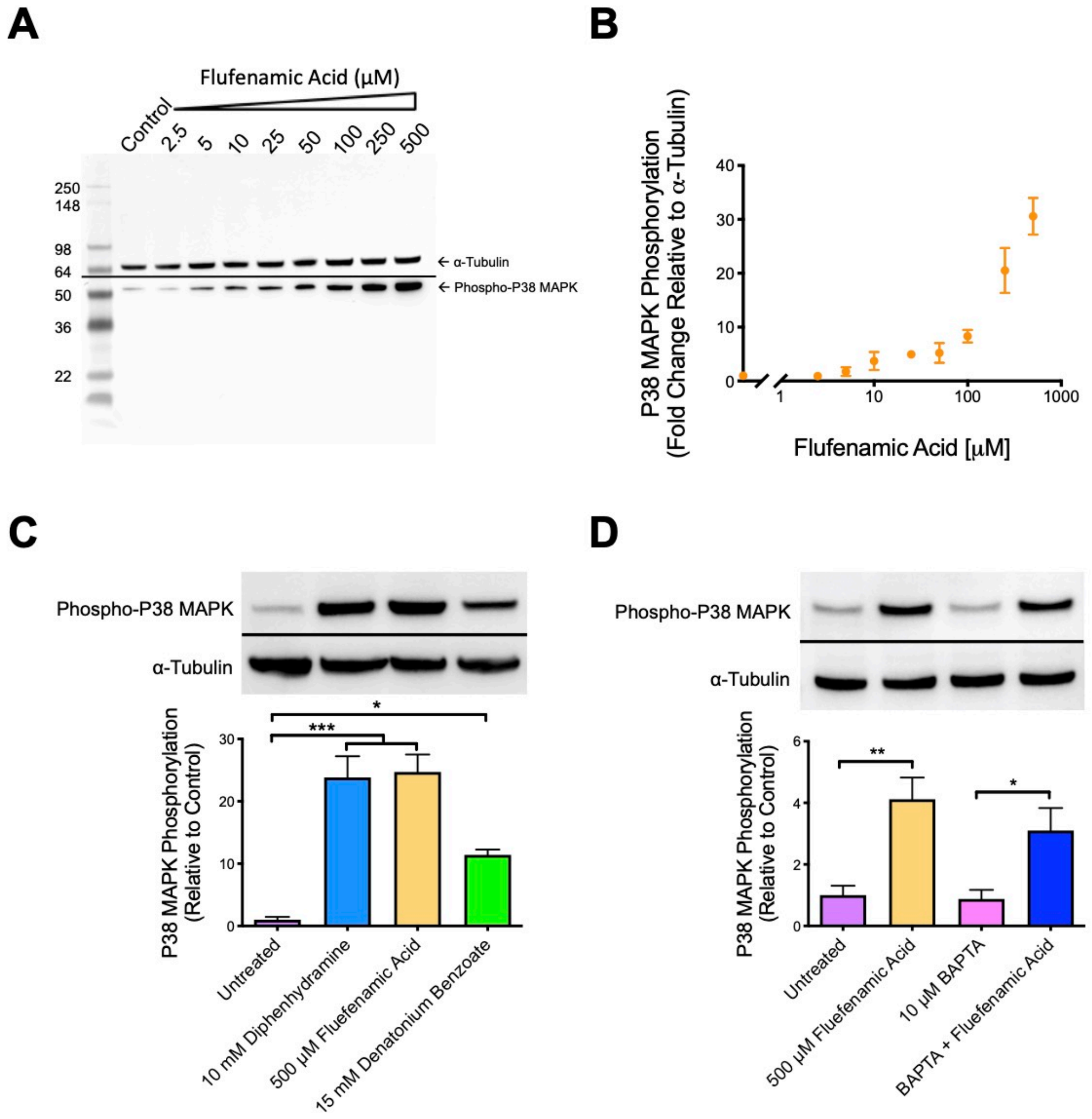


**Supplementary Fig. S16 Bitter compound-induced $Ca^{2+}$ release does not influence P38 MAPK phosphorylation. A** Western blot image depicting P38-MAPK phosphorylation relative to loading control α-tubulin (60 µg of protein per lane of protein gel). **(B)** Densitometry analysis of Beas-2B cells treated with either HBSS or 2.5-500 µM of flufenamic acid for 1 hour. Data points represent mean ± SEM of 3 immunoblot images. **(C)** Beas-2B cells treated with T2R agonists for 1 hour show an >10-fold increase in phospho-P38 MAPK via western blot. **(D)** BAPTA does not prevent P38-MAPK phosphorylation in Beas-2B cells pre-loaded with 10 µM BAPTA (1 hour) then treated with 500 µM flufenamic acid for 20 min. **(A,B)** Bar graphs showing the mean ± SEM of densitometry values of phospho-P38 MAPK relative to α-Tubulin signal combined from 3 experiments and analyzed by 1-way ANOVA with ***c*,** Tukey post hoc test; *$P$<0.05, ***$P$<0.001 or D) Bonferroni post hoc test; *$P$<0.05, **$P$<0.01.

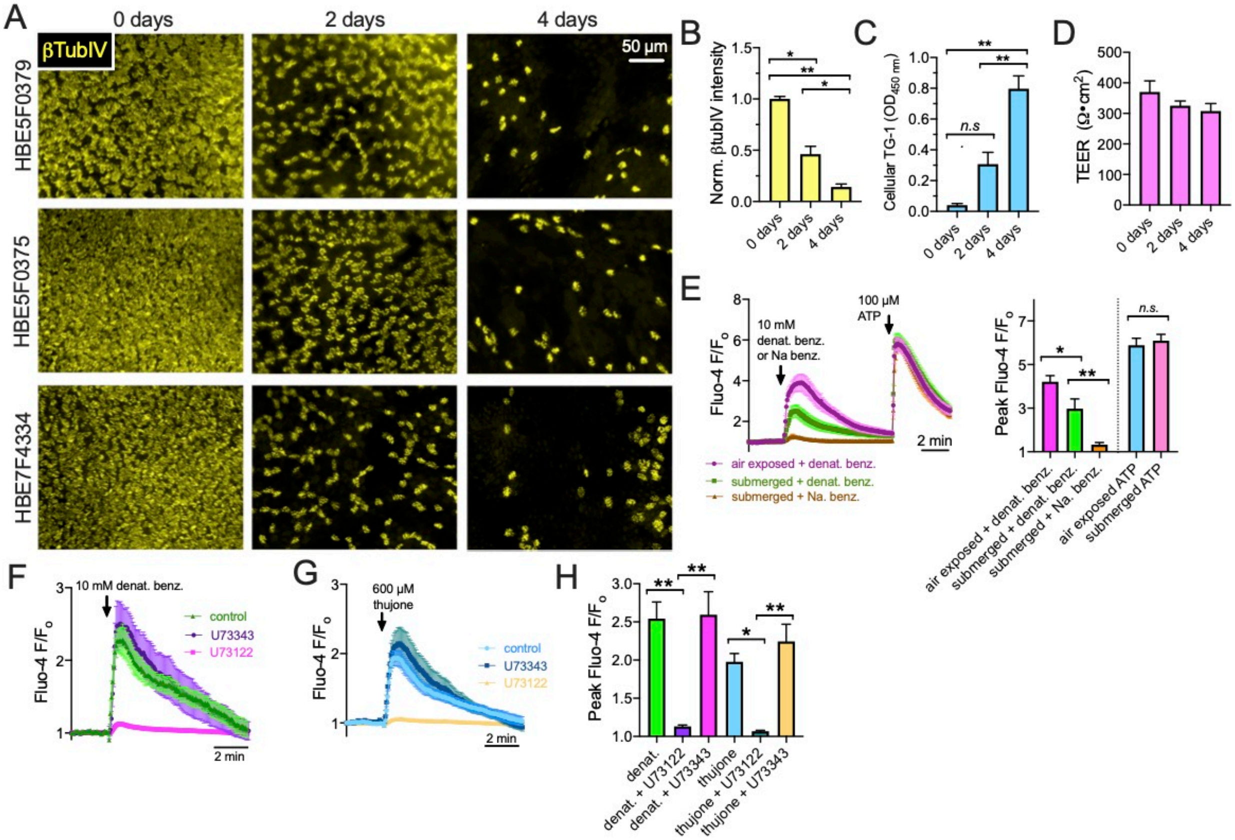


**Supplementary Fig. S17 De-ciliated bronchial epithelial cells retain $Ca^{2+}$ responses to bitter agonists.** It was previously shown that T2Rs in bronchial cilia detect denatonium benzoate [4, 40] to stimulate ciliary beating. Normal human bronchial epithelial (HBE) cells from 5 different donor lungs (Lonza, Walkerville MD) were cultured as previously described [2, 41] for 3 weeks at on transwells filters (Corning) at air liquid-interface in Lonza bronchial epithelial basal media (BEBM) supplemented with Lonza Singlequot™ supplements. After differentiation (day 0), the cells were subjected to 4 days of apical submersion, which induces squamous metaplasia and de-ciliation [42, 43]. **A** Immunofluorescence of cilia marker β-tubulin-IV (βTubIV; detected using mouse monoclonal antibody ONS.1A6, Abcam ab11315) showed marked loss of cilia as previously described [2]. De-identified donor numbers shown on the left, days of submersion shown on the top. Air-liquid interface cultures (ALIs) were fixed in 4% formaldehyde, blocked with 1% BSA and 1% normal donkey serum, then permeabilized with 1% triton and 0.2% saponin. Secondary antibody was conjugated to AlexaFluor488. Cells were visualized by with 10x objective on a wide-field microscope with GFP filters as described [2]. **B** Quantification of βTubIV immunofluorescence in cultures from 5 donors (1 ALI per donor at each time point showing decreasing intensity with 2 and 4 days apical submersion. Significance determined by one-way ANOVA with Bonferroni posttest; $*p<0.05$ and $**p<0.01$. **C** Cellular transglutaminase 1 (TG-1) expression was measured as a marker of squamous metaplasia was measured as described [2] using TG-1 ELISA (Aviva Systems Bio, San Diego, CA, Cat # OKCD01601). TG-1 expression went up with decreasing βTubIV. Significance determined by one-way ANOVA with Bonferroni posttest; $**p<0.01$ and *n.s.* = no significance difference. Bar graph shows mean ± SEM of data from 3 ALIs per time point from 3 separate donors (1 ALI per time point per donor, 9 ALIs total). **D** Transepithelial electrical resistance (TEER) was not altered by submersion, suggesting the epithelial barrier was intact. No significant difference by one-way ANOVA. Bar graph shows mean ± SEM of data from 4 ALIs per time point from 4 separate donors (4 ALI per time point per donor, 12 ALIs total). **E** $Ca^{2+}$ responses were measured as described [2] using Fluo-4 in 4-day submerged vs age-matched air-exposed cultures. Average traces mean ± SEM) from 6-9 ALIs from 3 donors (2-3 separate ALIs from each donor) per condition shown on the left and bar graph quantification of $Ca^{2+}$ peaks shown on the right. While the $Ca^{2+}$ response to denatonium benzoate was reduced, it was not eliminated by submersion. Response to purinergic agonist ATP was unchanged. There was no response to sodium benzoate in submerged cultures. Significance determined by 1-way ANOVA with Bonferroni posttest using paired comparison as indicated; $*p<0.05$, $**p<0.01$, and *n.s.* = not significantly different. **F-H** Responses to T2R agonists denatonium (representative traces in *f*) and thujone (representative traces in G) in submerged cultures were reduced by phospholipase C inhibitor U73122 (10 μM; 1 hr pre-treatment) but not inactive control U73343. Peak $Ca^{2+}$ responses summarized in bar graph shown in (H). Significance determined by 1-way ANOVA with Bonferroni posttest using paired comparison as indicated; $*p<0.05$ and $**p<0.01$.
